# Individual-level contextual alignment and aperiodic slope reveal improved comprehension of unexpected language

**DOI:** 10.1101/2025.10.28.685236

**Authors:** Ashley L. M. Platt

## Abstract

In everyday language use, individuals are required to comprehend complex, continuous and often unexpected linguistic inputs to effectively communicate with others. Evidence from other domains of processing such as memory and learning suggest that individuals these unexpected inputs can often be remembered or learnt just as well as predictable inputs in follow up testing. However, language comprehension research often focuses on how individuals process these stimuli rather than how the information which is gathered can be used at a later time. It is proposed that, in line with memory and learning, individuals can utilise unexpected language inputs to support a deeper understanding of the information which is being communicated. Further, this will be related to the interaction between an individual’s alignment to local context and inter-individual differences in information processing, such as aperiodic slope. The present study uses an alignment metric which measures the relationship between individual level N400 amplitudes (a neurophysiological metric of predictability) and surprisal (a text-collection-based metric of predictability), where higher alignment suggests the individual has a stronger preexisting schema. A steeper aperiodic slope has been suggested to reflect greater adaptation of predictive models in response to prediction errors; therefore, it is hypothesised that depth of language comprehension will be modulated by both alignment and aperiodic slope. To explore this, 30 participants (23F, 1 not reported; mean age: 22.2 years [SD:4.4]) listened to 12 short stories (3 genres, audio-visual presentation) while their electroencephalogram (EEG) was recorded. Comprehension was tested via 6 multiple choice questions per story. An additional combined analysis, including a reanalysis of 40 audio- modality participants, provided evidence across both modalities that comprehension was improved for more surprising stories. Further, when considering inter-individual differences, the relationship between predictability and comprehension was shown to be modulated by both alignment and aperiodic activity. Comprehension of unexpected stories (as measured by N400 amplitudes) was improved only lower aligned, steeper aperiodic slope individuals. This finding is considered a reflection of complementary processing strategies which are weighted towards incoming stimuli, increasing the likelihood of model updating and therefore resulting in improved comprehension of unexpected stories.

## 1 Introduction

In everyday language use, individuals are presented with complex, continuous and multimodal sensory input which the brain must seek to process in an efficient and effective way to best support real-time communication. It has been suggested that the brain achieves this efficient processing by making predictions about upcoming stimuli (Pickering & Garrod, 2007) and it is often concluded that the easier and more accurate these predictions are, the more efficient language comprehension is. This is supported by evidence from studies which focus on the real-time processing of linguistic information including increased reading and fixation times when processing unexpected words (e.g., Frank et al., 2013; Smith & Levy, 2013) and larger event-related potential (ERP) responses such as more pronounced N400 effects, thought to reflect increased processing effort (Kutas & Federmeier, 2011). The naturalistic language experience differs from these traditional laboratory settings, which often included controlled, isolated stimuli for the purpose of performing artificial tasks. This suggests that there may be more to learn about the full complexity of language by exploring processing in a naturalistic context (Alday et al., 2017; Brennan, 2016; Brennan & Hale, 2019; Zhang et al., 2021). One example of this complexity is in how individuals process unexpected information and how this impacts their overall understanding of the language content. Unexpectedness is currently often considered a hindrance to real-time language comprehension. However, this conclusion sits outside of other domains which rely on predictive perspectives, such as memory and learning, which commonly conclude that unexpected inputs can be remembered and learnt just as well (e.g., Quent et al., 2022) or better (e.g., Brod et al., 2018) than predictable inputs. The present study will build on a previous theoretical account of *Functional Unexpectedness*) and evidence for this provided by Platt (2025a) in the auditory modality. The aim is to extend this account to understand a multi-modal context and consider inter-individual differences in how individuals can make use of unexpected information.

### 1.1 Predictability outside of language comprehension

Current perspectives on the role of predictability in language processing suggest that the comprehension system operates most efficiently in conditions of higher predictability, but evidence from other processing domains suggests that there may be more complex relationships at play. In fact, in memory, it is commonly concluded that the presentation of unexpected stimuli can improve memory outcomes (Corley et al., 2007; Haeuser & Kray, 2023). Further, the process of learning is thought to be supported by mistakes, or errors in predictions, which allow individuals to adjust their internal expectations to integrate this new information. Memory and learning studies often use linguistic stimuli for experiments of this type and have demonstrated that when linguistic information is presented in an unexpected context (e.g., incongruent sentence continuations), individuals report better recognition memory for information encoding in these unexpected conditions (Haeuser & Kray, 2023). Extending this understanding to include a neurophysiological perspective, studies exploring the N400 component have also demonstrated how N400 amplitudes can reflect individual-level adaptation in the face of unexpected inputs. The N400 is an event related potential (ERP) component commonly researched in language and is broadly described as being sensitive to manipulations of expectations and subsequent fulfilment (Kutas & Federmeier, 2000, 2011). The N400 component is present for every word an individual processes and larger (more negative) amplitudes are associated with greater unexpectedness (Kutas & Federmeier, 2011). A study by Hodapp & Rabovsky (2021) proposes the functional significance of the N400 as an error-based implicit learning signal. They observed that individuals in a perceptual identification task had better implicit memory for words which had been presented in an unexpected context in a previous sentence reading task. As a result, they conclude that N400 amplitudes reflect adaptation to the current processing context and that this adaptation is driven by errors (i.e., unexpected input) (Hodapp & Rabovsky, 2021). Further, they note that this adaptation is modulated by N400 amplitudes as well, with larger N400 amplitudes providing a larger implicit memory benefit during the implicit memory task. Taken together, this perspective on the N400 component and evidence from memory studies which use linguistic stimuli, suggests that there is more understanding to be gathered regarding the role of predictability in language comprehension.

### 1.2 Functional Unexpectedness

In Platt (2025b), a framework was presented regarding the role of unexpectedness in language comprehension and how it might support comprehension depth. This perspective proposes that the functionality of unexpectedness is modulated by the presence of suitable predictive schemas at an individual level to support integration of new, unexpected information. It further suggests that the presence of these predictive conditions can be operationalised using an alignment metric, which is based on the relationship between N400 amplitudes and corpus-based lexical surprisal. Surprisal is defined as the negative log probability of a word given its previous context (Lowder et al., 2018; Venhuizen et al., 2019). Its relationship to language comprehension is well established, with studies indicating that higher surprisal is associated with longer reading times (Boston et al., 2008; Demberg & Keller, 2008; Smith & Levy, 2013) and larger N400 amplitudes in ERP research (Frank, 2015). Further, advances in predictive language models have provided an opportunity for large language models to expand the previous context component of surprisal calculations. As a result, it is suggested that word-by-word surprisal – as measured within a suitably large context window –can provide a measure for how individuals are expected to respond to language stimuli based on the probabilistic contingencies of the current context. By contrast, N400 amplitudes can reflect how they actually respond to the incoming linguistic stimuli. This assumption is supported by the correlation between N400 amplitudes and implicit memory demonstrated in Hodapp & Rabovsky (2021), providing a functional link between individual responses and behavioural outcomes. Platt (2025a) demonstrated that in a naturalistic story listening task, depth of language comprehension was increased for unexpected stimuli and that this was particularly evident in cases of lower alignment (where surprisal and N400 amplitudes were less related to each other). These results highlight that successful integration of unexpected information is related to alignment to local context but that it may be conditions where alignment is lower that integration is more likely. It is suggested that this finding could be related to an individual’s likelihood to rely on either their existing predictive models or on the unexpected incoming information. In the case of lower alignment, individuals may weight their model updating towards the incoming unexpected stimuli and update models, accordingly, allowing for better integration and subsequent comprehension. This perspective is in line with the Bayesian brain hypothesis (or Bayesian inference) which outlines that beliefs (which can be equated to existing predictive models) are updated based on the precision (i.e., the width of the probability distribution of the information) of both the belief and the incoming sensory input (Adams et al., 2013; Feldman & Friston, 2010). A belief held with high precision has a narrow probability distribution highly concentrated over the most likely expectation (Adams et al., 2013). Prediction errors act as evidence (or lack thereof) for the prior belief and act to shift the posterior belief towards the prior or sensory evidence in proportion to their relative precision (Adams et al., 2013). The role of precision in language processing has been explored in relation to the N400 component, which was proposed in Bornkessel-Schlesewsky & Schlesewsky (2019) as a precision-weighted prediction error signal which modulates internal model updating. Support for this perspective was provided by a study of individual-level processing strategies which further suggested that the precision weighting of incoming information and existing models may differ between individuals (Bornkessel-Schlesewsky, Sharrad, et al., 2022). This study compared N400 responses to speakers who differed in their likelihood to use a canonical adjective order (e.g., huge grey elephant) or a non-canonical order (e.g., grey huge elephant). Individuals rapidly adapted their prediction models to reflect the intra-experimental conditions, with prediction reduced for the less likely order, regardless of global expectations which are shifted towards the canonical order (Bornkessel-Schlesewsky, Sharrad, et al., 2022). Further this adaptation was strongest for individuals with a steep aperiodic slope and lower individual alpha frequency (IAF). Building on this finding, it is proposed that individuals who are less aligned to story-level context have less precision over their prior beliefs (i.e., internal predictive models) and therefore bias their posterior beliefs, relied on in a comprehension task, towards the sensory input. These conclusions highlight an important consideration for the interplay between internal and external predictive conditions in understanding the role of predictability in naturalistic language comprehension. Therefore, to further explore this perspective, the current study will present stimuli in a multimodal context and include a metric of interindividual differences in information processing (as measured by aperiodic activity).

### 1.3 External processing conditions and the role of multimodal stimuli

Considering the role of multimodal stimuli in naturalistic research is important, as in everyday communication language comprehension often occurs in both an audio and visual modality and as a result is accompanied by a range of non-linguistic features. Further, psycholinguistic research demonstrates that language processing occurs via the combination of multiple information sources to form an interpretation (for a discussion see Onnis & Spivey, 2012) . In fact, eye movement and ERP research has demonstrated that non-linguistic cues play an important role in supporting real-time comprehension (Knoeferle, 2015). Despite this, language studies often utilise single modality stimuli such as only audio or visual presentation (e.g., Dambacher et al., 2006; Russo et al., 2020). When multimodal stimuli are incorporated, they are often included to create an artificial discrepancy between the two streams of information. For example, studies which examined incongruency between gestures and spoken information, such as Wu & Coulson (2005), reveal larger N400 amplitudes in incongruent gesture and speech conditions. These studies provide a foundation for understanding the language comprehension process and provide an opportunity to extend this to naturalistic stimuli where artificial manipulations are not present.

Additionally, a common focus for multimodal explorations of language processing is processing of speech in adverse listening conditions. In such studies, audio stimuli are presented in clear and degraded conditions to explore how visual information can support processing in these conditions. Visual information, provided by gestures and visible speech, has been shown to enhance language comprehension under adverse listening conditions by providing phonological information about the speech signal (Drijvers et al., 2019; Ross et al., 2007). A study by Drijvers et al. (2019) found that listeners allocate their visual attention to the face, in comparison to hand gestures, when speech was degraded and reason that listeners do so to extract more information to aid comprehension. As such, multimodal language processing studies suggest that viewing a speaker’s articulartory movement provides complementary information to enhance comprehension, especially in noisy processing conditions (Ross et al., 2007). In the context of processing unexpected information and in light of findings from Platt (2025a) , these conclusions suggest that audio-visual processing of unexpected stimuli may support subsequent comprehension by shifting prescision weighting towards the sensory signal. Under circumstance where processing is weighted towards the existing model (i.e., in response to unexpected information), additional visual information may provide external conditions which make functional unexpectedness more likely.Further, the increased precision of incoming external information as provided by an audio-visual input may also shift this processing towards stimuli, in a similar way to lower alignment conditions. In suggesting that external processing conditions may influence the precision-weighting of model updating, it is important to also consider differences in internal processing conditions.

### 1.4 Interindividual differences in internal language processing conditions

When exploring how individuals process unexpected information it is essential to consider how this is related to their unique, internal processing conditions. At a baseline, individuals differ in how they process the world around them which can lead to differences in various domains of processing such as attention, speech production and processing and working memory capacity (Broadway & Engle, 2011; Daneman & Green, 1986; Huettig & Janse, 2016; Just & Carpenter, 1992; Kidd et al., 2018). In the processing of language, researchers have attempted to control for this variability using group level attributes such as age (DeLong et al., 2012; Maquate et al., 2025; Maquate & Knoeferle, 2021) or gender (Hyde & Linn, 1988; Özçalışkan & Goldin-Meadow, 2010). However, various accounts have begun to demonstrate that electrophysiological metrics can account for a considerable amount of interindividual differences. This is an important consideration for an account which seeks to explain how individuals can support comprehension of unexpected information and what characteristics may make this functionality more likely.

One interindividual metric in neurophysiological research which has been proposed to explain differences in internal processing conditions is aperiodic activity. Aperiodic activity has been discussed in relation to individual differences in cognitive ability and demonstrated to modulate prediction-related N400 effects (Dave et al., 2018; Hill et al., 2022). Aperiodic activity adheres to a 1/f power distribution, in which spectral power decreases with increasing frequency, and is often referred to as 1/f activity (Hill et al., 2022). The slope component of this relationship has been suggested to change dynamically and is related to factors including age and task in addition to an individual’s cognitive state (Donoghue et al., 2020; He, 2014; Voytek et al., 2015). Further, studies have demonstrated that individuals with a steeper slope have more pronounced prediction-related N400 effects (Dave et al., 2018) and adapt their internal models more readily to novel input (Bornkessel-Schlesewsky, Sharrad, et al., 2022). In an artificial language learning experiment, Cross et al., (2022) provided evidence that learning of certain grammatical rules was predicted by variability in 1/f slope. Additionally, Bornkessel-Schlesewsky, Sharrad, et al. (2022) also found that 1/f slope was related to an individual’s alignment to speaker-based grammatical expectations. They conclude that that improved prediction, as evidenced by steeper 1/f slope, may relate to an individual’s ability to flexibly adapt their predictive coding models to a local context. Aperiodic (1/f) slope has also been suggested to correlated with amount of neural noise in the brain, with a steeper slope representing more synchronised neural firing and therefore less neural noise (Pertermann et al., 2019). Increased neural noise (reflected in aperiodic slope) has also been associated with neural gain (Bensmann et al., 2018; Buckley & Toyoizumi, 2018; Servan-Schreiber et al., 1990), a principle of information processing in the brain which can amplify a systems response to incoming stimuli and may support individuals in establishing higher precision predictive models. Taken together, these studies suggest that when exploring an individual’s capacity to utilise unexpected information to support language comprehension may be modulated by interindividual differences in processing. It is proposed here that the interplay between alignment and aperiodic slope will determine the precision weighting of incoming signals and determine how individuals process novel information.

### 1.5 The present study

The current study aimed to explore the role of internal and external processing conditions in modulating the relationship between predictability and language comprehension. Using the same naturalistic stories as Platt (2025a), now presented in the audio-visual modality, individuals viewed the stories while their electroencephalogram (EEG) was recorded. Comprehension was measured by a series of multiple-choice questions, designed to test an individual’s contextual understanding of the story content. Story-level alignment to local context was measured by the relationship between individual-level N400 amplitudes and surprisal. Further, to explore the role of aperiodic activity in these relationships, a reanalysis of existing data from Platt (2025a) was combined with the current data to provide a more suitable sample size for examining inter-individual differences. In the audio-visual modality it is predicted that lower alignment will support greater comprehension of unexpected stories, in line with findings from Platt (2025a). Additionally, a steeper aperiodic slope will also be associated with greater comprehension depth due to weighting of internal processing conditions towards incoming stimuli. Insights from the present study will outline the internal and external processing conditions which modulate the relationship between predictability and language comprehension.

## 2 Methods

### 2.1 Participants

The audio-visual study sample consisted of 30 participants (23F; 1 not reported) who were healthy, right-handed adults between the ages of 18-40 years (mean: 22.2, SD: 4.4). Handedness was assessed using the Flinders Handedness Survey (Nicholls et al., 2013), a brief self-report measure of skilled handedness. Additional participant inclusion criteria included no consumption of recreational drugs in the last 6 months or be taking any regular medication which would affect the EEG. Participants also had normal or corrected vision, reported normal auditory acuity and did not have any diagnosed psychiatric, neurological, cognitive or language disorders and to avoid language confounds, participants were native English speakers who had not learned a second language before the age of 5 years. The combined analysis included reanalysis (Platt, 2025a) of an additional 40 participants (27 female, mean age: 24.6[SD: 5.94]) who fit the same criteria as above but who were allocated into an audio-only condition. The combined sample therefore included 70 participants (50F, 1 not reported, mean age: 22.2 years [SD: 4.4]). The experiment protocol was approved by the University of South Australia’s Human Research Ethics Committee (protocol number: 204945).

### 2.2 Materials and measures

#### Audio-visual stimuli

The story stimuli for this experiment included 12 short stories classified as fiction, newspaper and science (4 stories per genre) with an approximate duration of 3-4 minutes. Stories were recorded in mono-speaker format by a native speaker of Australian English. Stories were presented in audio-visual format, where audio-visual recordings included the speaker from shoulders up. These stories were the same recordings as used in Platt (2025a) , where they were presented as audio only Several characteristics per story were calculated for use in the statistical analysis. This included word position, word duration, sentence length, word frequency, idea density, word audio frequency and intensity. Story segmentation, to calculate word position, word duration and sentence length, was completed via WebMAUS Basic (Schiel, 1999).Unigram frequency values were computed from the Open Subtitles corpus for English (751 million words) as made available by van Paridon & Thompson (2021).Computerised Propositional Idea Density Rater (CPIDR; Brown et al., 2008) was used to calculated ID for each short story. Idea density (ID; also known as Propositional Density or P-Density. Kintsch & Keenan, 1973) utilises written or oral text samples to calculate the number of ideas expressed relative to the number of words used and has linked to linguistic ability (Farias et al., 2012), language production (Kemper et al., 2001) and comprehension (Kintsch & Keenan, 1973). Acoustic characteristics were calculated using Praat version 6.4.04 (Boersma & Weenink, 2024). The current study used this software to calculate word duration, fundamental frequency and intensity. The descriptive characteristics of each story are outlined in Table 1.

**Table 1:** Characteristics for audio-visual stimuli at genre level [M (SD)]

| Genre | Mean duration | Mean word count | Mean Idea density | Mean Word frequency | Mean Word duration | Mean Sentence length | Mean Fundamental frequency | Mean Intensity | Mean Surprisal |
| --- | --- | --- | --- | --- | --- | --- | --- | --- | --- |
| Fiction | 2m50s | 430.0<br>(42.1) | 0.52<br>(0.03) | 5014689.5<br>(589760.1) | 0.30<br>(0.03) | 14.41<br>(5.51) | 227.6<br>(6.96) | 58.53<br>(0.55) | 5.30<br>(0.77) |
| Newspaper | 2m57s | 433.5<br>(11.5) | 0.46<br>(0.04) | 5161020.6<br>(707570.0) | 0.33<br>(0.01) | 26.30<br>(3.43) | 227.9<br>(1.84) | 57.88<br>(0.54) | 5.73<br>(0.65) |
| Science | 3m10s | 446.8<br>(57.8) | 0.52<br>(0.05) | 4465183.5<br>(637829.1) | 0.35<br>(0.01) | 19.85<br>(1.74) | 225.6<br>(5.86) | 57.29<br>(1.44) | 5.17<br>(0.24) |

#### Aperiodic (1/f) activity

Participants’ eyes-closed resting state EEG recordings were used to calculate aperiodic slope estimates. Aperiodic (1/f) intercept and slope estimates were calculated using the YASA toolbox (Vallat & Walker, 2021) in Python. YASA utilises the irregular-resampling auto-spectral analysis (IRASA) method for separating oscillatory and aperiodic activity (Wen & Liu, 2016). By-participant intercept and slope estimates were computed as means of pre and post resting-state recordings from electrodes; Fz, F3, F4, F7, FC5, FC1, C3, CP5, CP1, Pz, P3, P7, O1, Oz, O2, P4, P8, CP6, CP2, Cz, C4, T8, FC6, FC2, F4, F8.

#### Comprehension probes

Comprehension probes were designed to measure a story-level semantic understanding and were presented at the end of each story. Six multiple choice questions were presented for each story (72 total) and were designed by the primary researcher. Questions were designed by the primary researcher based on an approach outlined in Delgado & Salmerón (2021) and Ozuru et al. (2007). These studies suggest that comprehensions questions can be designed to probe a contextual understanding of stimuli and not only rely on recall memory. As such, the final questions were chosen to reflect if listeners had gained an in-depth understanding of the stories, beyond the surface level information provided by text. Comprehension probes were piloted for clarity and to ensure ceiling effects were not present. Depth of comprehension for statistical modelling, as indicated by comprehension probes, was calculated at story level to reflect the proportion of correct answers per story (i.e., a score ranging from 0-1). See Figure 1 for example of comprehension questions.

**Figure 1:**
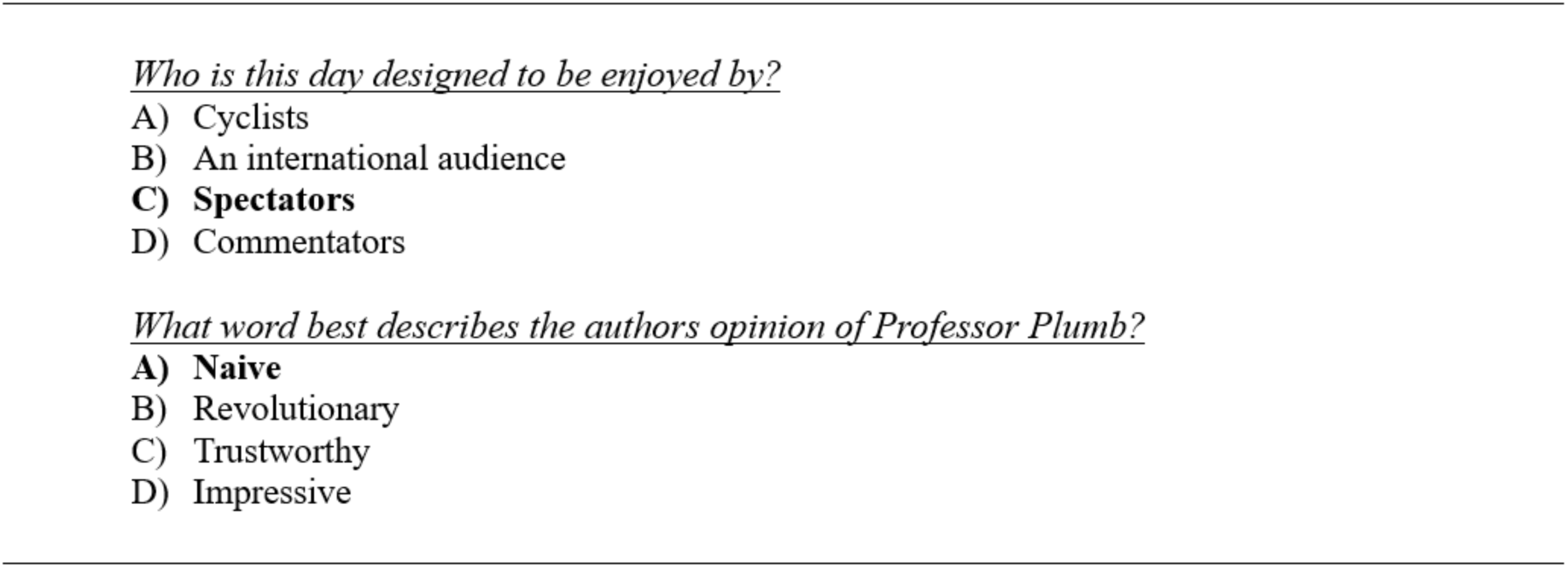
*Example of multiple choice comprehension probes. Bold indicates correct answer*.

#### Surprisal calculation

Surprisal was calculated at word level and included a context window of all words prior to the current word. Thus, the amount of contextual information used for the surprisal calculation increased over the course of each story. Word-level surprisal values were used in the alignment models and were then averaged at story-level for the comprehension models. Surprisal is the negative log probability of a word given its preceding context (Lowder et al., 2018). The equation to calculate is as follows:

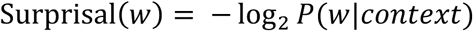

To increase the context window of our surprisal calculation, we used GPT-2 (Generative Pre-trained Transformer-2), which is a pretrained large-scale transformer model trained on billons of words of text with the objective to predict the next word (Radford et al., 2018, 2019). GPT-2 was utilised via functions *GPTConfig, GPT2Tokenizer, GPT2LMHeadModel and GPT2Model* from the *transformers* package in Python version 3.10.14. Occasionally GPT-2 calculations split up words into ‘tokens’. In these cases, we summed the value of the tokens to provide the whole word surprisal, as outlined in Stanojević et al. (2023).

#### Electroencephalography (EEG) recording and pre-processing

EEG was recorded with a 32-channel Brainvision ActiCHamp system (Brain Products GmbH, Gilching, Germany) with a sampling rate of 500 Hz. Electrodes were placed according to the 10-20 system. Vertical and horizontal eye movements were monitored with two electrooculogram (EOG) electrodes placed beside the outer canthus of the right and left eyes, as well as two additional EOG electrodes slightly above and below the left eye. The online reference and ground were AFz and FPz, respectively. Impedances were kept below 10k Ω. EEG pre-processing and analysis was carried out in MNE Python version 1.6.1 (Gramfort et al., 2013). EEG signals were re-referenced to the average of left and right mastoids (TP9 & TP10). Independent Component Analysis (ICA) was utilised to correct EOG artifacts. Independent components found to correlate most strongly with EOG events (via the *create_eog_epochs* function in MNE) were excluded. A band-pass filter of 0.3-20Hz was applied to the raw data to remove signal drift and high frequency noise. All following analyses are based on epochs from -0.2-1 seconds relative to the onset of each word in each story.

### 2.3 Procedure

Eligible participants attended a 2 hour in-lab testing session which included EEG setup and testing in addition to collection of resting state and idea density metrics. The story task ran for roughly 50 minutes and required participants to listen to each story and then answer the six associated depth of comprehension questions. Participants were initially presented with a welcome screen, followed by a brief instructions screen before being prompted to ‘press any key to begin’. Each story was preceded by a 10 second count down before a video of the speaker was presented for the duration of each story. Following story completion, participants were presented with instructions followed by a series of comprehension questions and instructed that they could take this time to relax before continuing with the story task. The interim instruction screen and comprehension questions were not assigned with a maximal reaction time, and the experiment would only progress following participant engagement. Story recordings were played though a loudspeaker at a comfortable volume for the participant. Story order presentation was determined by assignment to one of four randomly generated story orders in which stories of the same genre never appeared more than twice in a row. List presentation was counterbalanced across participants.

### 2.4 Data analysis

The present study involved two separate analyses. The first included the audio-visual sample (N=30) and involved three major stages: an initial identification of a region of interest, alignment modelling and comprehension modelling. The second analysis used the ROI and alignment values from analysis 1, combined with the values produced in a previous analysis of the audio sample (N=40) (Platt, 2025a) to explore the relationship between depth of comprehension, alignment and 1/f (aperiodic) slope (final sample size, N = 70). The analysis pipeline is outlined in Figure 2.

**Figure 2:**
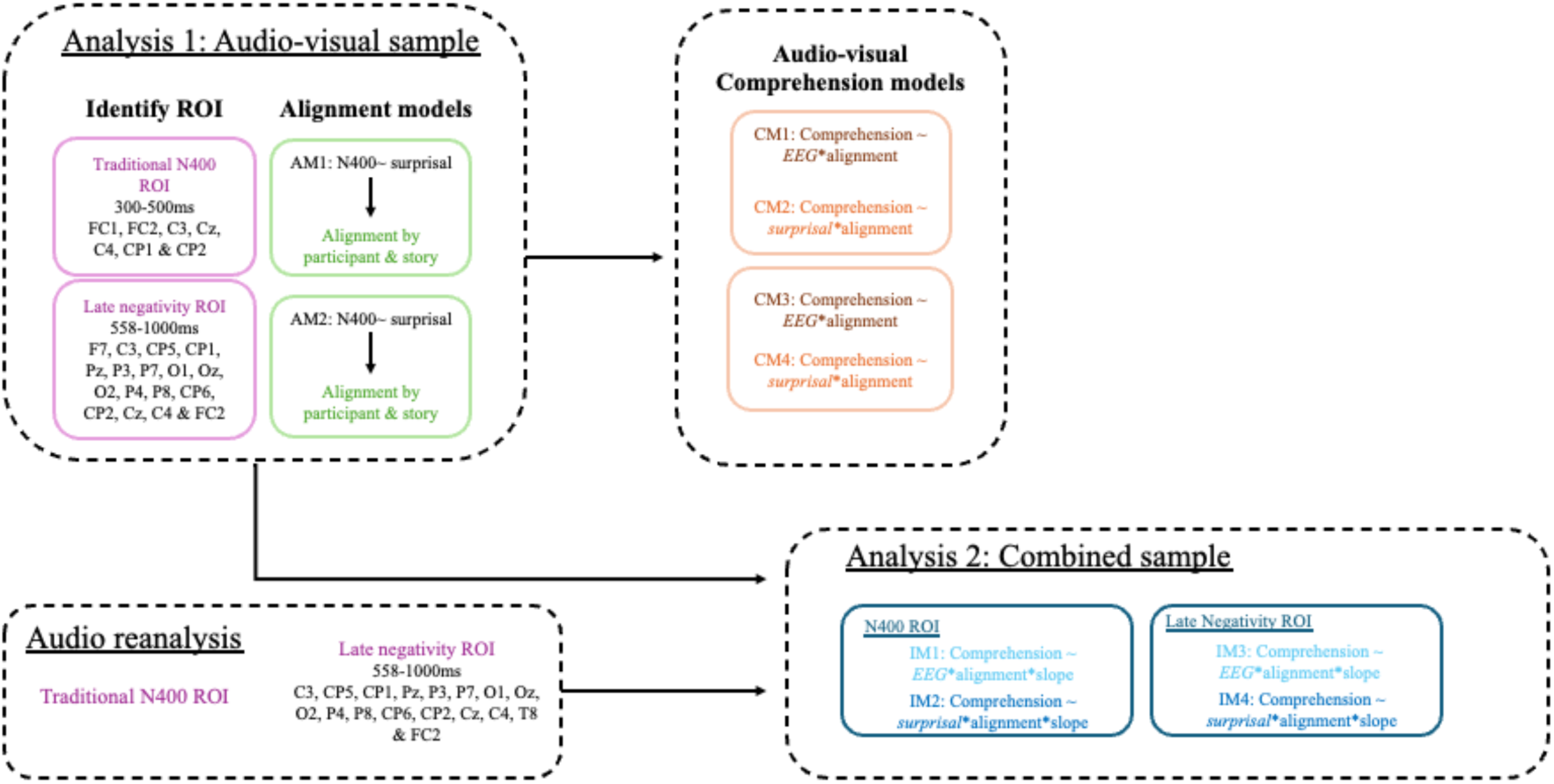
*Schematic of analysis pipeline Analysis 1: Audio-visual only*

The audio-visual sample was analysed using the pipeline outlined in Platt (2025a) namely a whole-head single trial analysis to identify a data-driven ROI followed by alignment and comprehension models. Regression-based ERPs (rERPs; Smith & Kutas, 2015) were calculated for all content words in order to examine the predictors of interest (surprisal and EEG amplitudes) while controlling for continuous covariates of no interest. These covariates included pre-stimulus activity (-200-0 ms prior to word onset; Alday, 2019), word-position in the story, word duration, log-transformed word frequency, average fundamental frequency and intensity. A group analysis was then conducted per time point and electrode to identify ROIs where EEG amplitudes significantly correlated with surprisal. The identified ROI and a traditional N400 ROI (300-500ms) were then used to compute alignment models 1 and 2. This analysis again mirrored the alignment models in Platt (2025a), and therefore the models (AM1 & 2) included fixed effects of surprisal, log unigram frequency, word position and prestimulus activity and their interactions. Control predictors were also consistent with Platt (2025a) and included word duration (previous & current), log-transformed previous word frequency, average fundamental frequency and intensity. The random effects structure for each model was determined using a parsimonious model selection procedure to avoid over-parameterisation (Bates et al., 2021; Matuschek et al., 2017), with final models including random intercepts by subject, story, word and channel. Final model structure is outlined below and was consistent between the two ROIs.

***Alignment model structure:*** average n400 amplitudes ∼ pre-stimulus activity * word position * log frequency * story-level surprisal + duration + average fundamental frequency + average intensity + previous word frequency + previous word duration + (1 + pre-stimulus activity + log frequency + word position | subj) + (1 + story-level surprisal | subject & story) + (1 + pre-stimulus activity + word position + story-level surprisal | word) + (1 | channel)

As a measure of individual- and story-level alignment, the beta values from each alignment model were then used to predict comprehension depth. The comprehension models (CM1, 2,3,4) were computed for each ROI and key predictor (EEG amplitudes & surprisal) and included a quadratic surprisal or EEG predictor to account for non-linear relationships. Story-level control predictors included average sentence length, average word duration, average log transformed word frequency, idea density and genre. The random effects structures, as determined by parsimonious model selection, were consistent across all four models and included random slopes by participant and story.

#### Analysis 2: Combined audio and audio-visual with individual differences

To enable the exploration of individual differences with a larger sample size, data from an audio only condition of the same experiment (Platt, 2025a) was combined with the audio- visual sample. Both the audio and audio-visual whole-head analysis identified comparable clusters. Therefore, the combined analysis will refer to these ROIs together as a ‘late negativity’ ROI. Comprehension models were then computed per ROI (N400 or late negativity), key predictor (EEG amplitude or surprisal) and aperiodic slope. Consistent with the audio-visual analysis above, each model included story level control predictors of average sentence length, average word duration, average log transformed word frequency, idea density and genre. The random effects structure for models IM1, 2 & 4 included a random intercept by condition | subject and a random slope by story. IM3 only included random slopes by subject and story.

## 3 Results

### 3.1 Behavioural outcomes

Table 2 outlines participant comprehension scores by condition. Across all genres, the average comprehension score was 70.5%. All participants’ total comprehension scores were above 50% indicating sufficient engagement with story stimuli. There were two cases in the audio-visual condition in which comprehension scores for one story were removed due to recording system errors. In these cases, individual percentages have been adjusted to account for a difference in total questions asked.

**Table 2:** *Comprehension scores by genre for audio-visual sample*

| Condition | Genre | Number of<br>observations | Mean % (SD) |
| --- | --- | --- | --- |
| Audio-visual | All stories | 358 | 70.5 (10.4) |
|  | Fiction | 118 | 72.5 (11.8) |
|  | Newspaper | 120 | 71.5 (11.7) |
|  | Science | 120 | 67.5 (13.6) |

### 3.2 Frequency effect ERP

As a sanity check to determine whether the current data showed expected modulation of N400 amplitude by unigram frequency we computed grand average low and high frequency content words. The N400 component is known to be modulated by word frequency with higher N400 amplitudes for low versus high frequency words (Kutas & Federmeier, 2011). This frequency effect is visualised in Figure 3 with a peak N400 amplitudes at roughly 700ms post word onset. This is later than the expected peak of 400ms but may reflect the ongoing processing of a continuous linguistic signal which can vary by word frequency. A similar latency was present in a study by Alday et al. (2017) who also used naturalistic stimuli. This finding was also consistent with the frequency effect presented in Platt (2025a), which had a similar shifted latency and peaking around 700ms post word onset.

**Figure 3:**
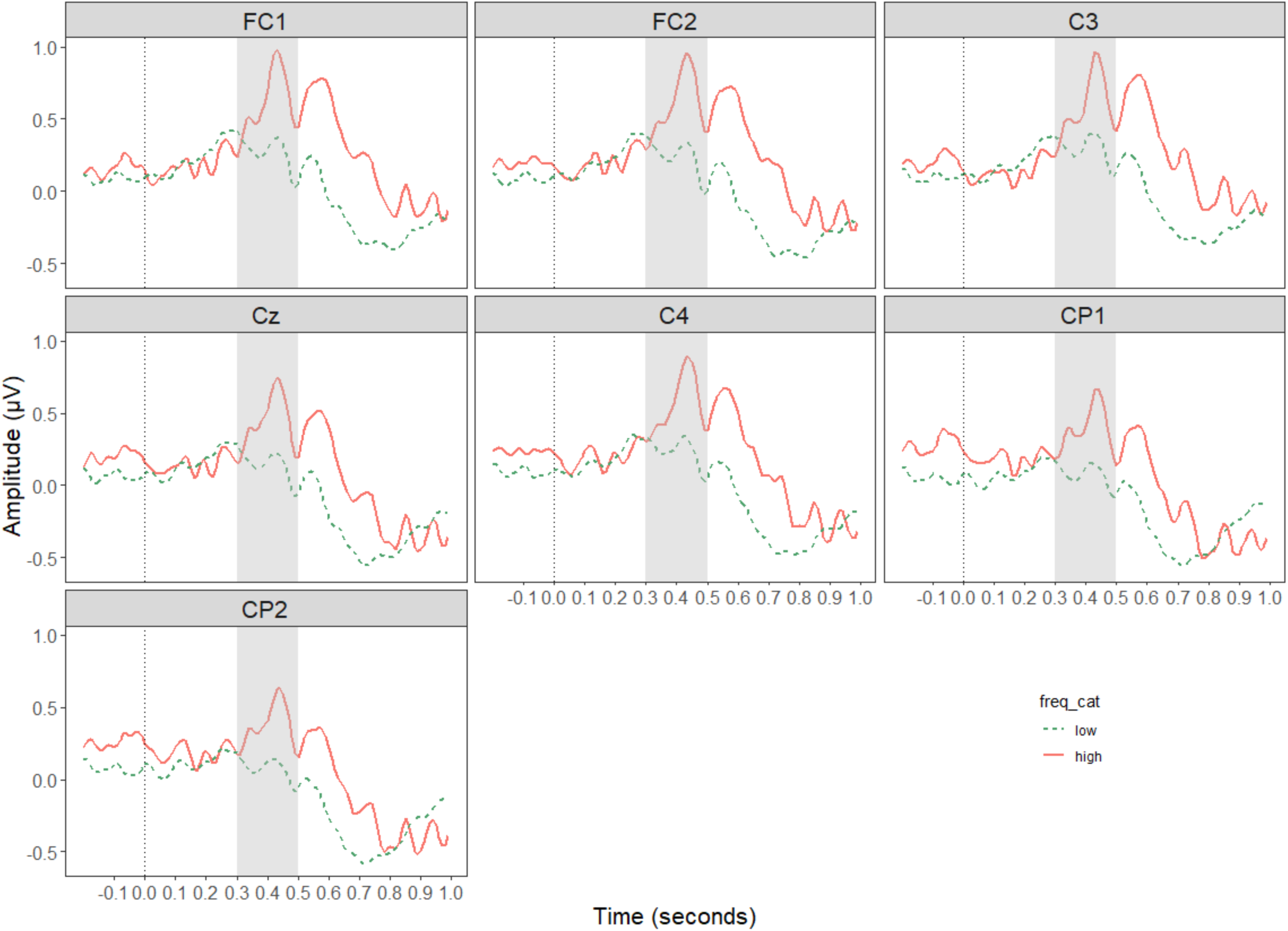
*Grand average ERP plots of frequency effect. Lines are separated by word frequency and amplitudes are plotted with negativity downwards. Highlighted region identifies traditional N400 ERP time window of 300-500ms*.

**Figure 4:**
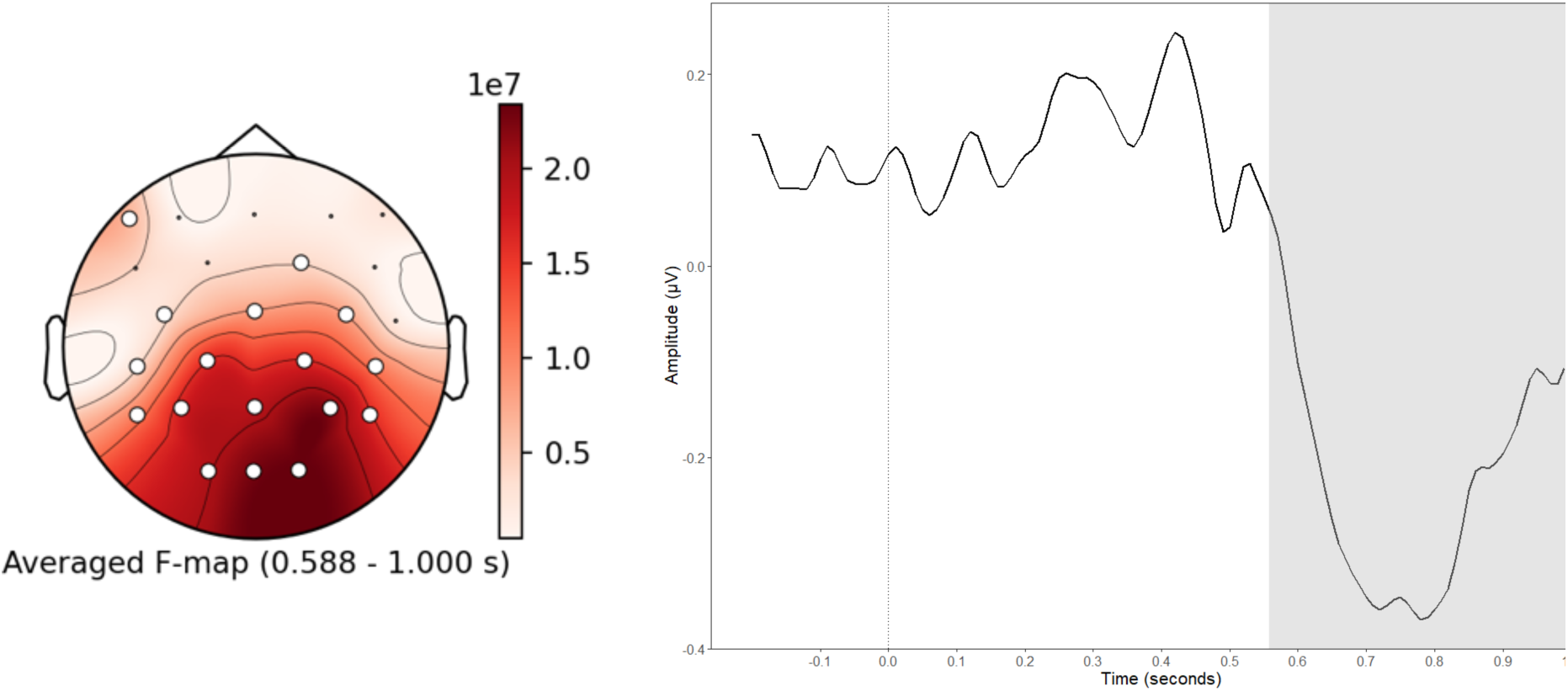
*Spatio-temporal characteristics and grand average plot for audio-visual late negativity cluster*.

### 3.3 Analysis 1: Audio-visual analysis

#### Alignment models

The whole-head single trial analysis identified one region of interest in which the relationship between surprisal and EEG amplitudes was significant. This cluster ranged from 558-1000ms and encompassed electrodes F7, C3, CP5, CP1, Pz, P3, P7, O1, Oz, O2, P4, P8, CP6, CP2, Cz, C4 and FC2 and presented as a late negativity (588-1000ms). See Figure.4 for the spatio-temporal characteristics of the cluster.

A mixed effects model was computed for each region of interest to predict EEG amplitudes from story-level surprisal to determine individual-level alignment to local context. The traditional N400 window (AM1) showed a significant highest order interaction between word position, log transformed word frequency and story-level surprisal (Estimate = 0.05, Std. Error = 0.02, z = 2.37, p = 0.02). This interaction effect is visualised in Figure 5. See Supplementary Material B for model outputs.

**Figure 5:**
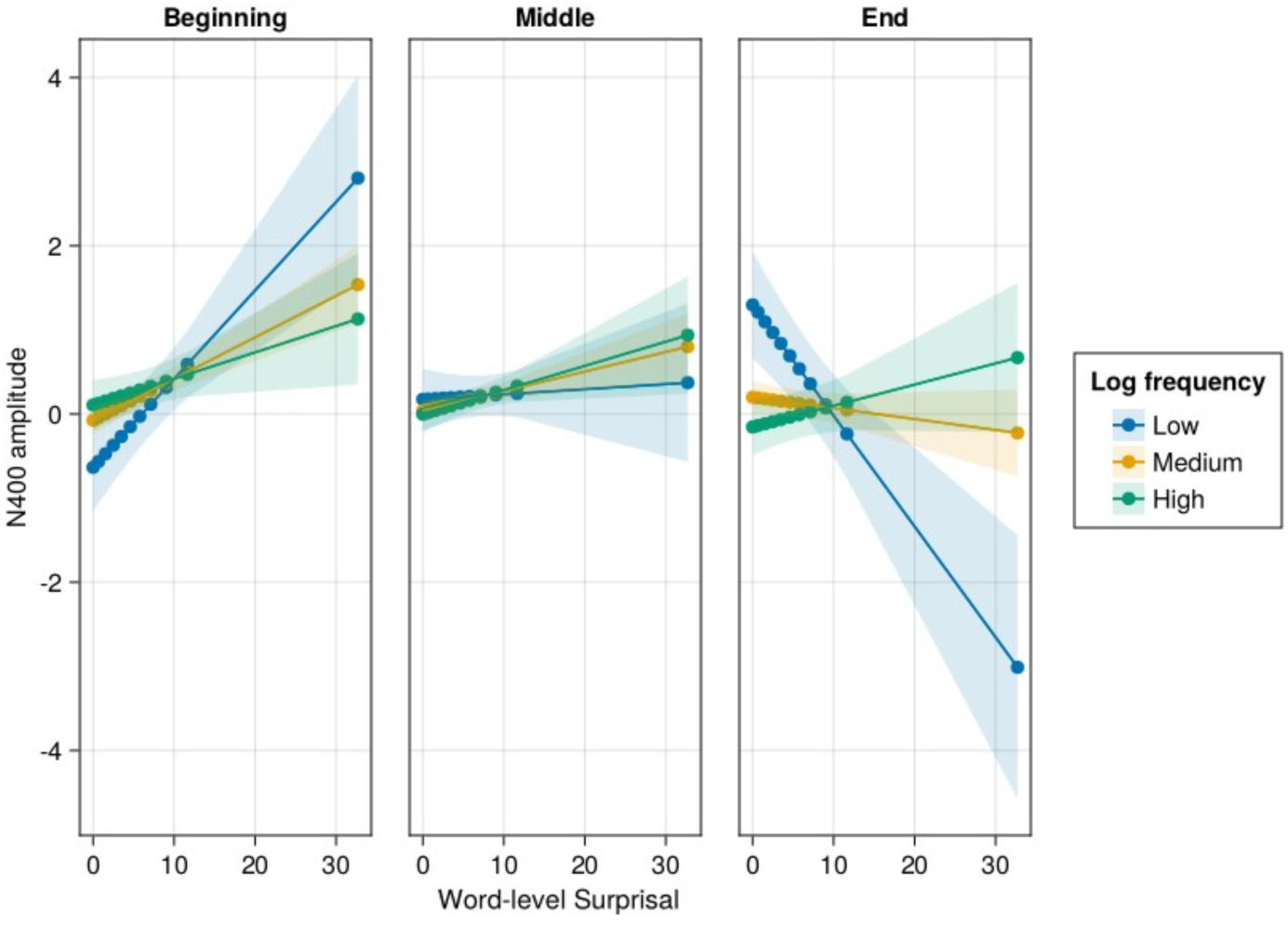
*Interaction between N400 amplitude and story-level surprisal by word frequency and position in story. The x-axis is surprisal, whilst the y-axis represents fitted values of N400 amplitude. The graph is faceted by story position and low, medium, and high frequency words are represented in separate slopes across each facet. Filled colours are 95% confidence intervals. The late negativity ROI (AM2) showed a significant interaction of word position and story-level surprisal (Estimate = -0.07, Std. Error = 0.01, z = -5.63, p <.001). This interaction effect is visualised in* Figure 6.

**Figure 6:**
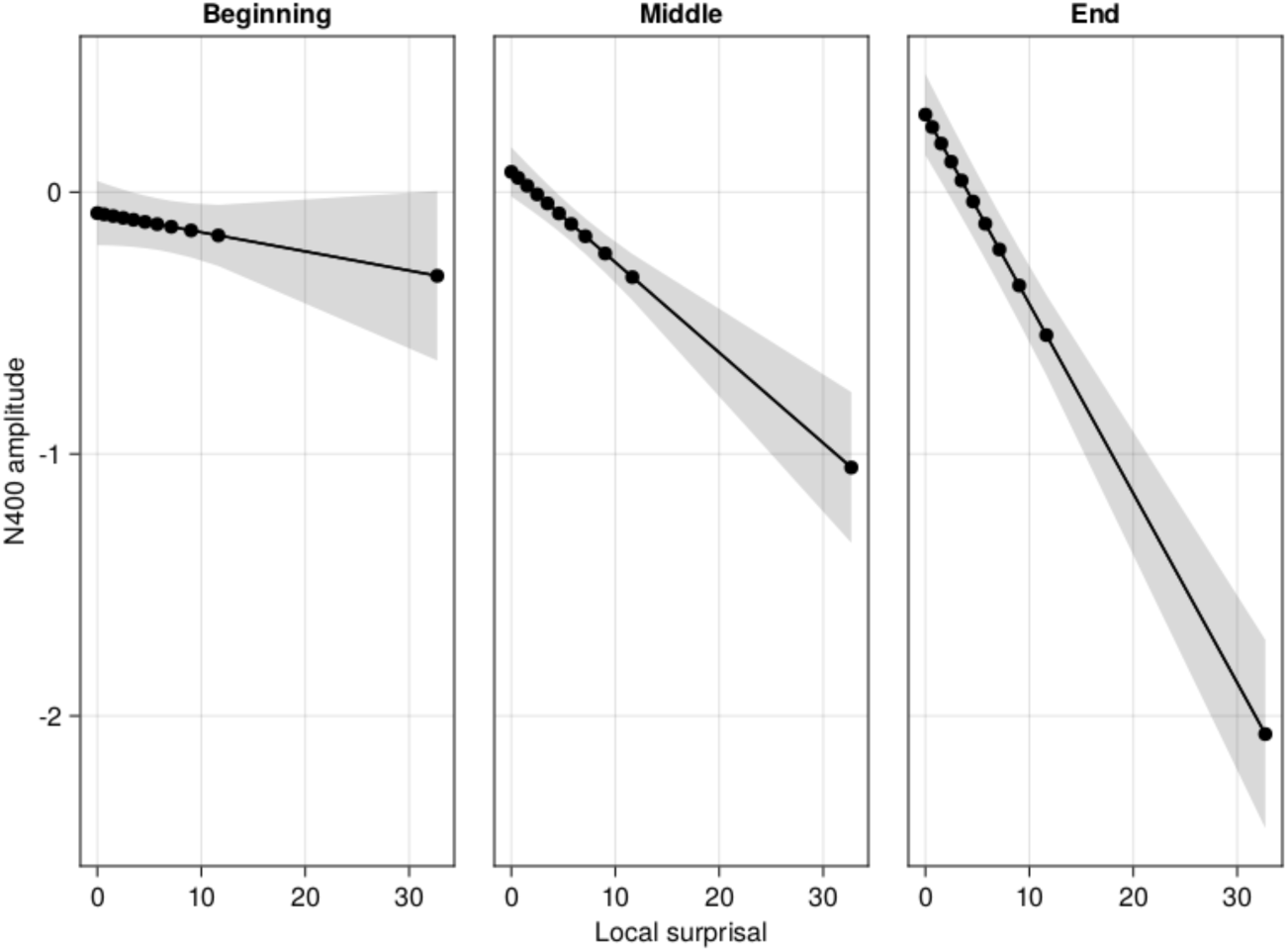
*Interaction between N400 amplitudes and story-level surprisal by word position. The x-axis is surprisal, whilst the y-axis represents fitted values of N400 amplitude. The graph is faceted by story position and filled areas are 95% confidence intervals*.

The random slopes of story-level surprisal by story and participant from these alignment models are utilised in the following comprehension models. The distribution of these values for each ROI is displayed in Figure 7 and the variation of alignment values by genre is displayed in

**Figure 7:**
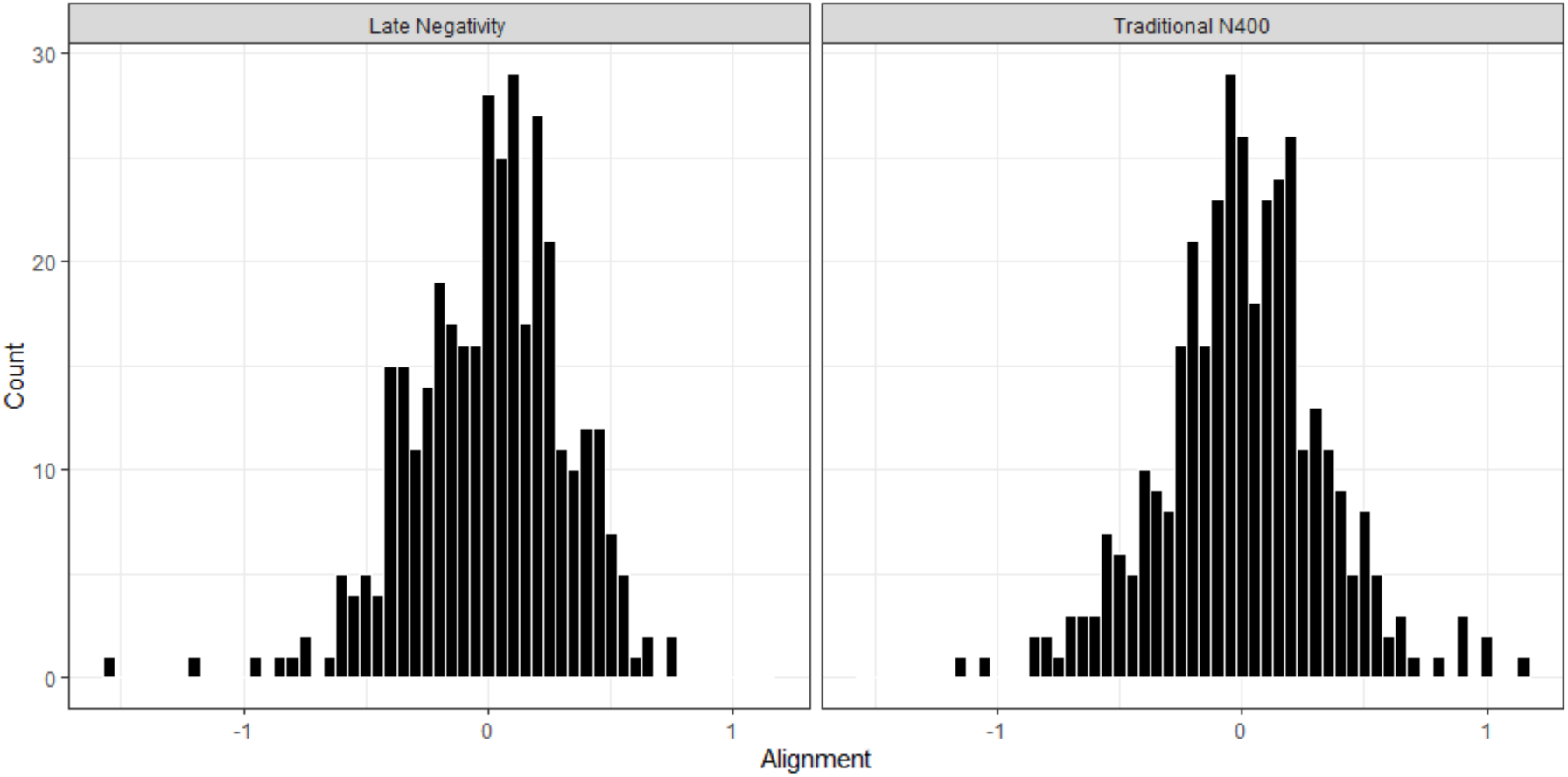
*Distribution of alignment values for Whole-head analysis ROI (LHS) and Traditional N400 ROI (RHS)*

**Figure 8:**
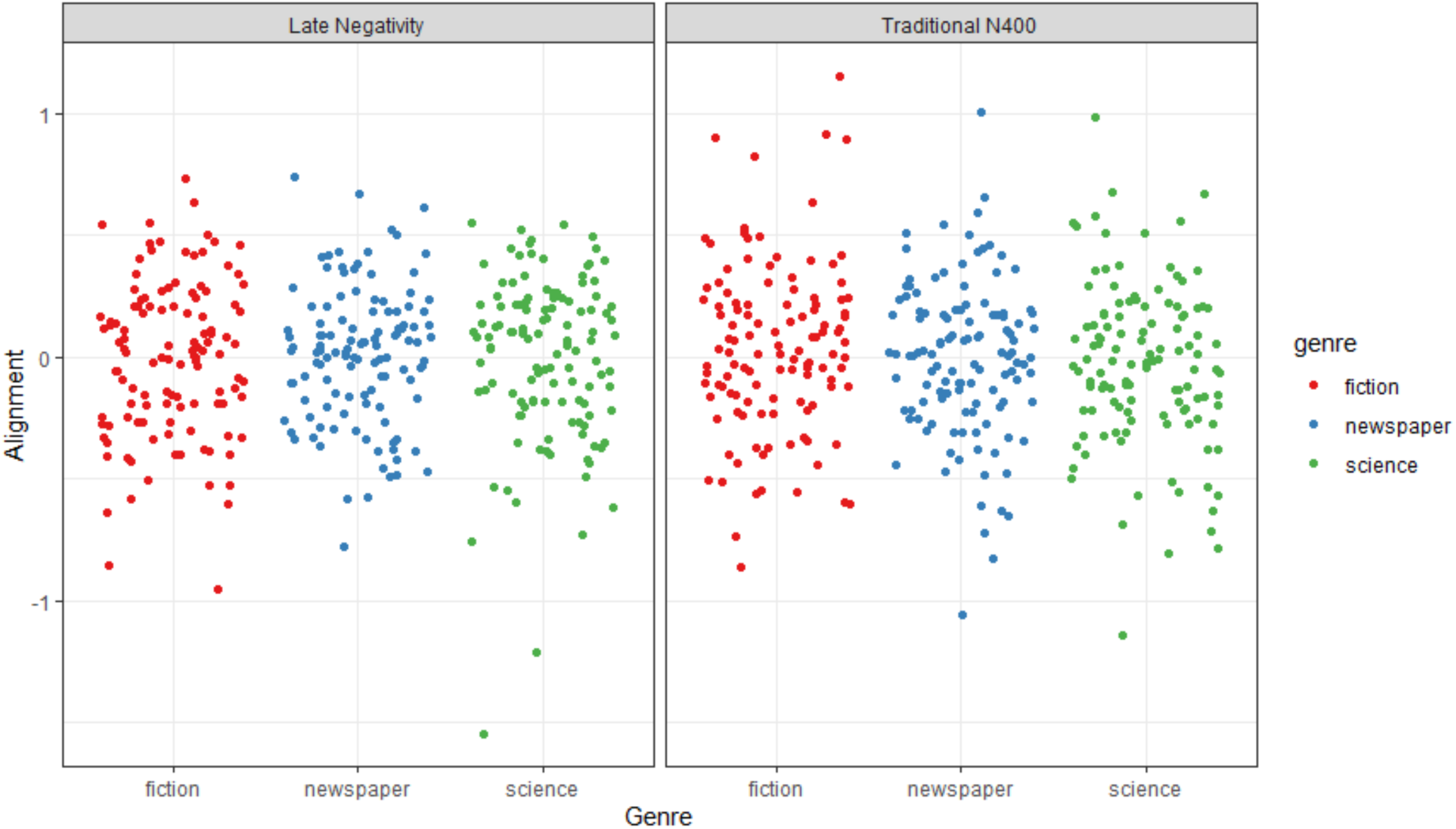
*Alignment variation by genre for Whole-head analysis ROI (LHS) and Traditional N400 ROI (RHS)*

#### Comprehension models

A comprehension linear mixed effects model per ROI (traditional N400 or late negativity) and each predictability metric (average N400 amplitudes or average surprisal) was computed. All models showed main effects of control variables average sentence length, average word duration and genre. The surprisal models further showed an effect of average word frequency and idea density. See Supplementary Material B for full model outputs.

#### EEG models (CM1&3)

LMMs for each ROI using story-average EEG amplitudes showed no further effects than the control variables.

#### Surprisal models (CM2&4)

Both surprisal models showed significant main effects of surprisal and surprisal^2^ and an interaction of surprisal and surprisal^2^ (equivalent to surprisal^3^) . Model results are reported in Table 3. Figure 9 visualises the main effect of surprisal in the traditional N400 ROI, with both models indicating that comprehension scores increase in line with increased surprisal.

**Figure 9:**
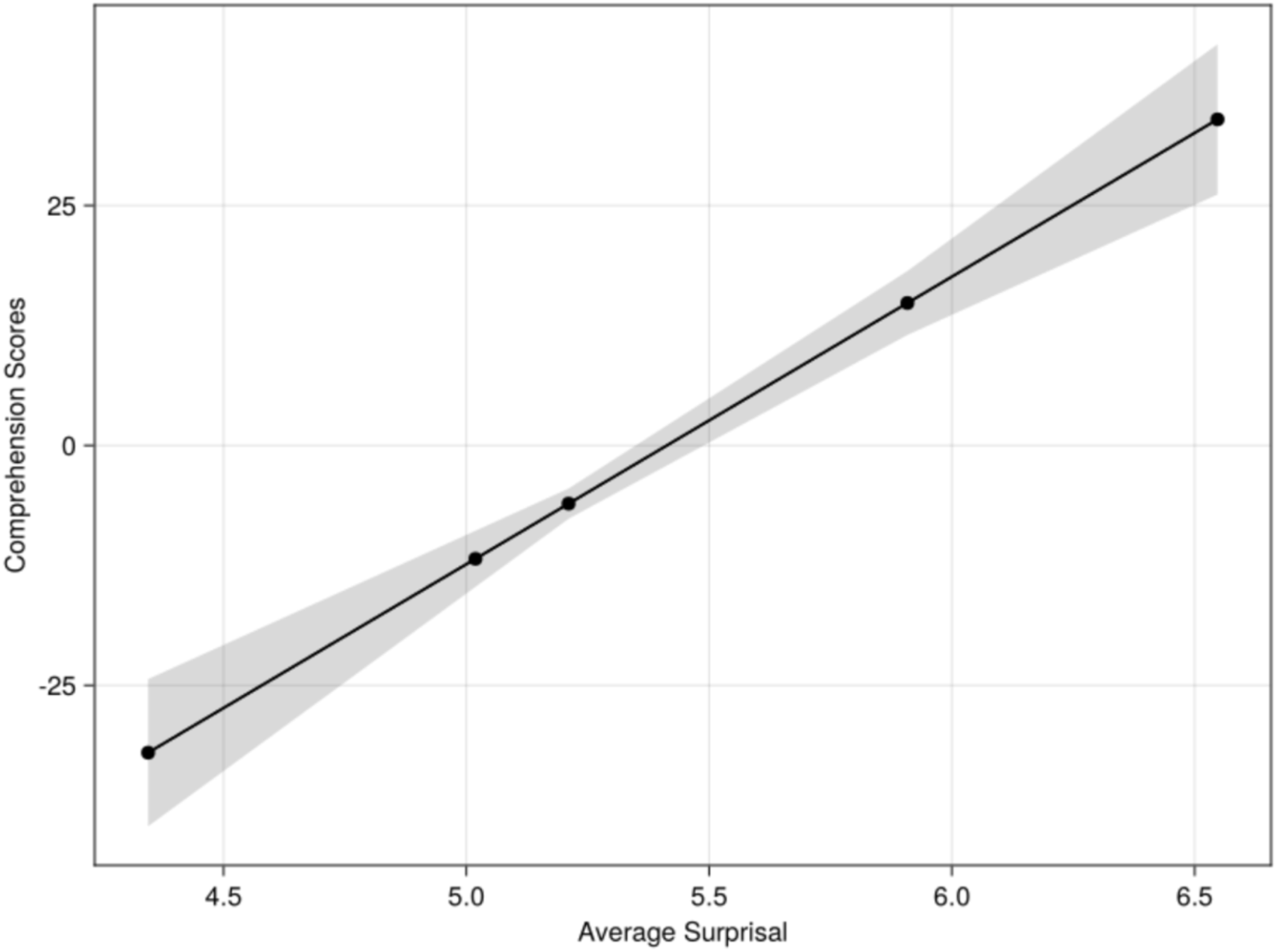
*Significant main effect of surprisal in predicting comprehension scores for the traditional N400 analysis ROI surprisal model using the audio-visual data set*.

**Table 3:**
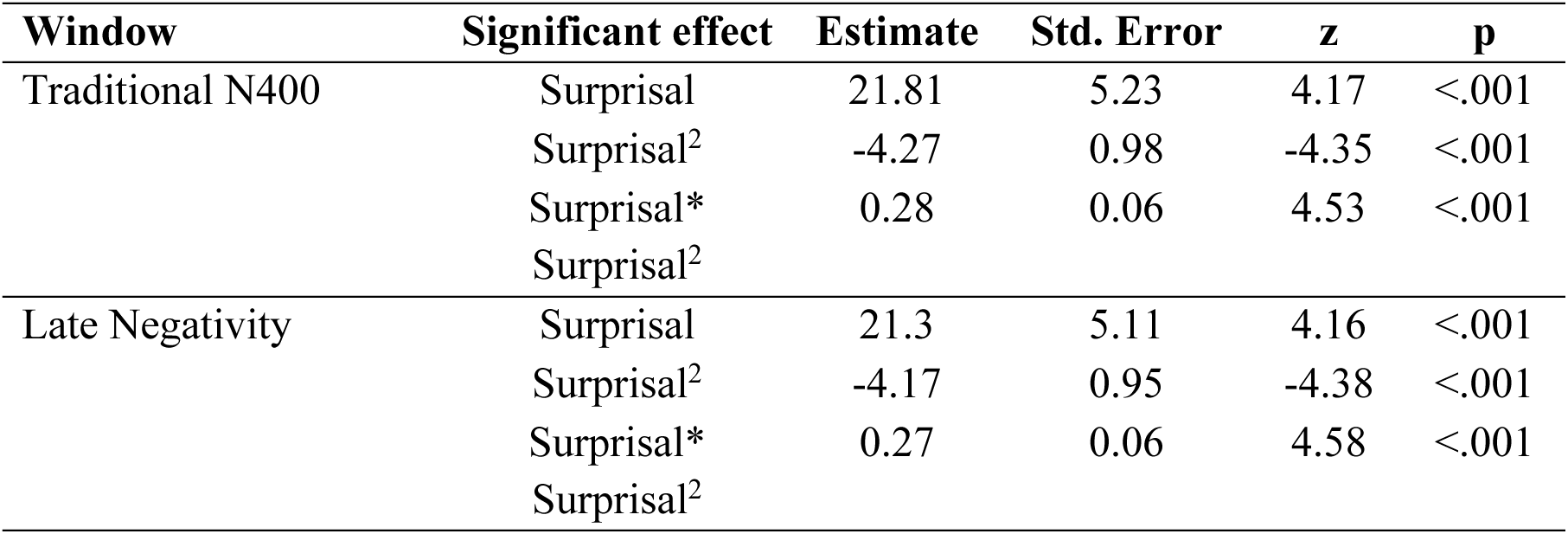
*Summary of significant effects for audio-visual analysis surprisal models*.

### 3.4 Analysis 2: Individual differences models

A comprehension linear mixed effects model per ROI (traditional N400 or late negativity) and each predictability metric (average N400 amplitudes or average surprisal) was computed using the combined data set (reanalysis of audio and audio-visual samples) and including aperiodic slope as a predictor. All models showed main effects of control variables average sentence length, average word duration, idea density and genre. The surprisal models further reported an effect of average word frequency. Modality was included as a covariate but did not interact with the other key predictors and will not be interpreted. See Supplementary Material B for full model outputs.

#### EEG predictor models

The traditional N400 EEG model (IM1) showed a significant main effect of alignment (Estimate = -0.46, Std. Error = 0.16, z = -2.92, p = .004) in which lower alignment was associated with greater comprehension scores (see Figure 10). An additional main effect of average N400 amplitudes was found (Estimate = -0.37, Std. Error = 0.14, z = -2.58, p = .01) where smaller (more positive) N400 amplitudes were associated with better comprehension (Estimate = -0.37, Std. Error = 0.14, z = -2.58, p = .01) (see Figure 11) A higher order interaction effect between aperiodic slope, alignment and average N400 amplitudes was also found (Estimate = 0.65, Std. Error = 0.23, z = 2.89, p = .004). This effect is visualised in Figure 12 and suggests that the relationship between average N400 amplitudes and comprehension depth is modulated by aperiodic slope and alignment. A functional relationship (greater N400 amplitudes associated with better comprehension) is found for lower story-level alignment and steeper aperiodic slope.

**Figure 10:**
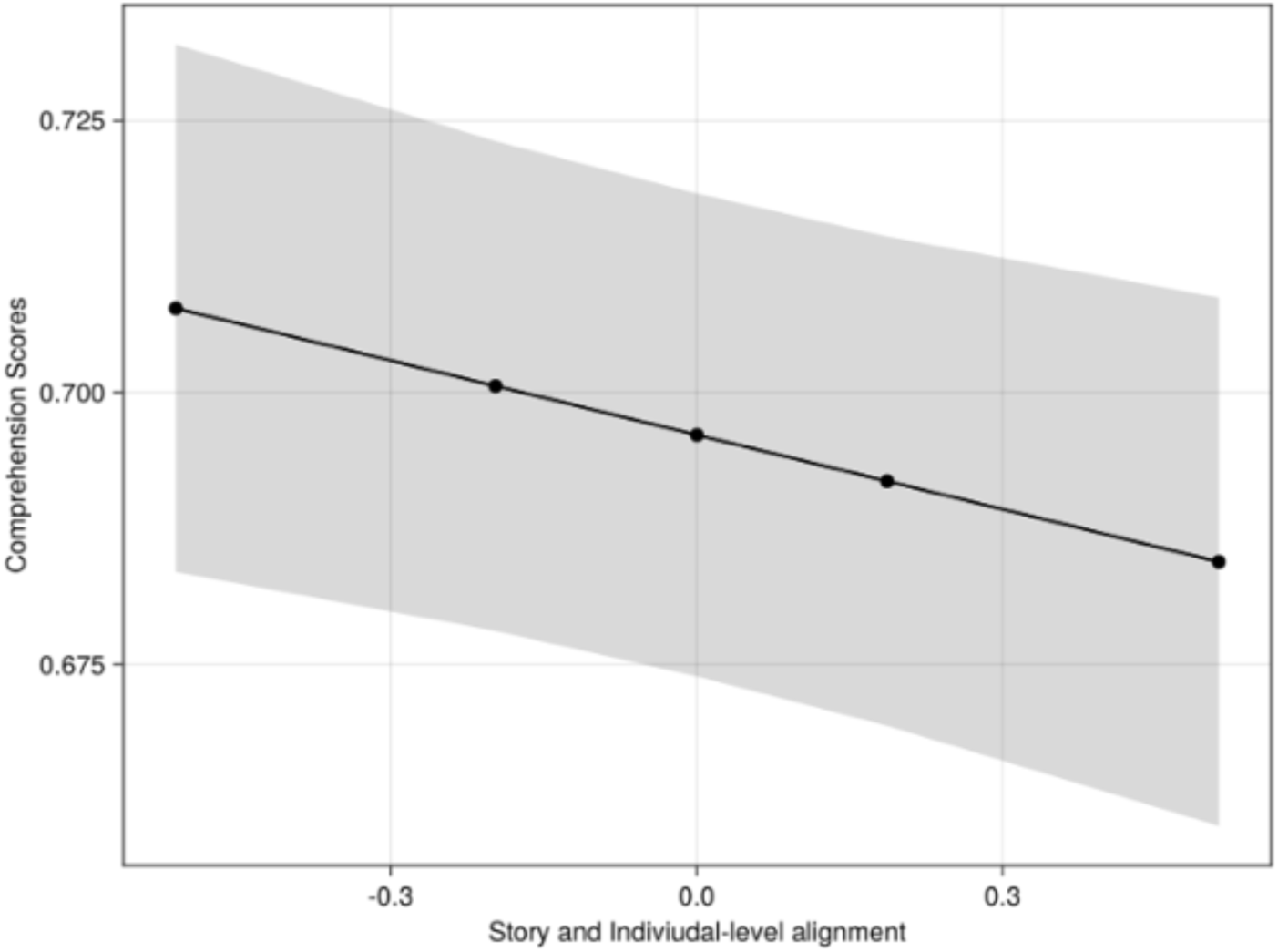
*Main effect of story and individual-level alignment from IM1. The x-axis is alignment, and the y-axis is fitted comprehension scores. The shaded area represents a 95% confidence interval*.

**Figure 11:**
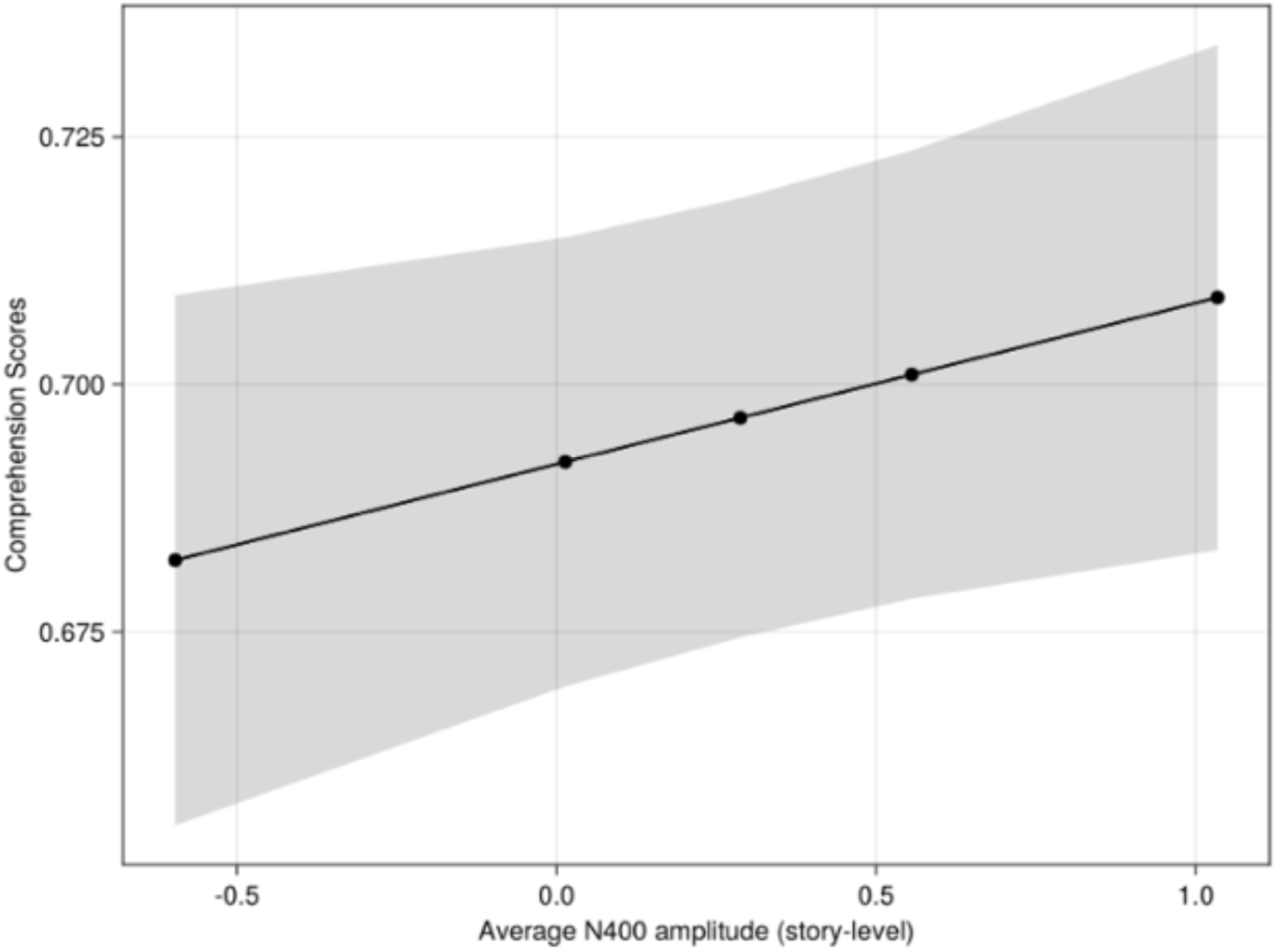
*Main effect of average N400 amplitudes from IM1. The x-axis is alignment, and the y-axis is fitted comprehension scores. The shaded area represents a 95% confidence interval*.

**Figure 12:**
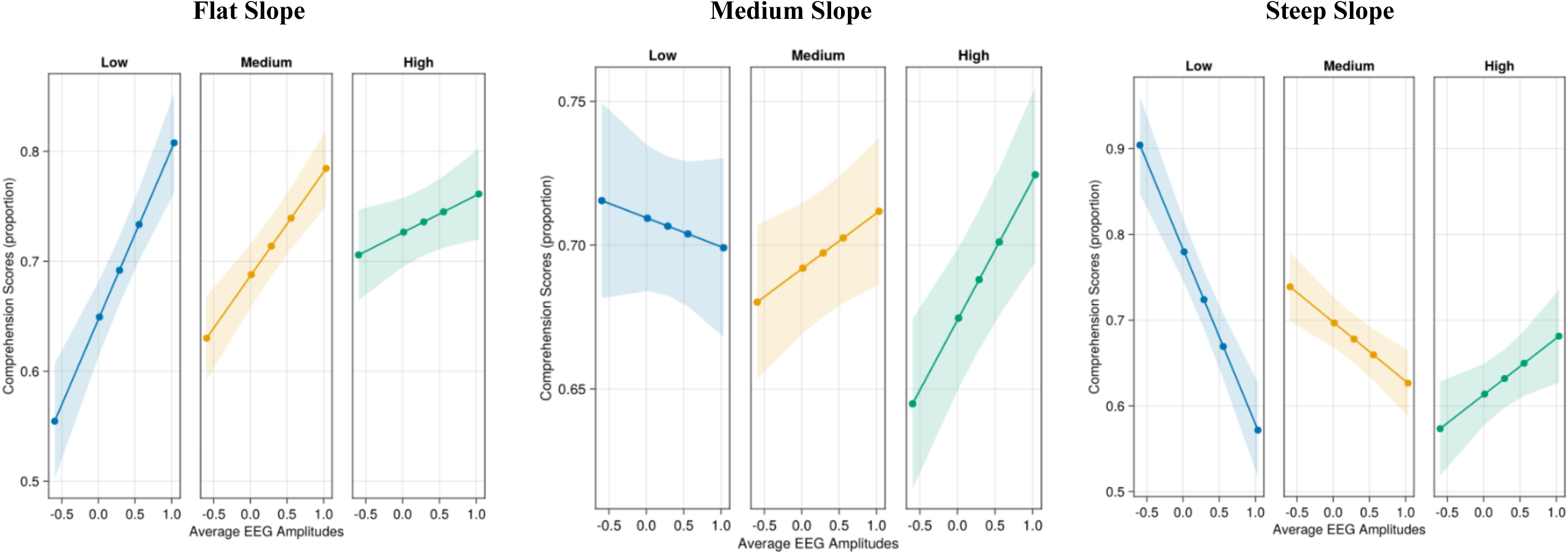
*Plot of highest order interaction between of aperiodic slope, alignment and average EEG amplitudes for each level of slope from IM1. The x-axis is average EEG amplitudes, and the y-axis is fitted comprehension scores. The plots are trichotomized for visualisation purposes for three levels of alignment and slope. Alignment is further visualised by colour with low, medium and high, alignment represented by blue, yellow and green respectively*.

The late negativity EEG model (IM3) showed a significant main effect of alignment (Estimate = -0.46, Std. Error = 0.17, z = -2.79, p = .005) and consistent with IM1, lower alignment was associated with higher comprehension scores. An additional main effect of quadratic average N400 amplitudes was also present in this model (Estimate = 0.54, Std. Error = 0.25, z = 2.15, p = .032) and the positive estimate suggests a U-shaped relationship between N400 amplitudes and comprehension. There was also a significant interaction effect of aperiodic slope and alignment (Estimate = -0.33, Std. Error = 0.12, z = -2.81, p = .005). Figure 13 visualises this effect and indicates that a steeper slope and lower story-level alignment are related to greater comprehension.

**Figure 13:**
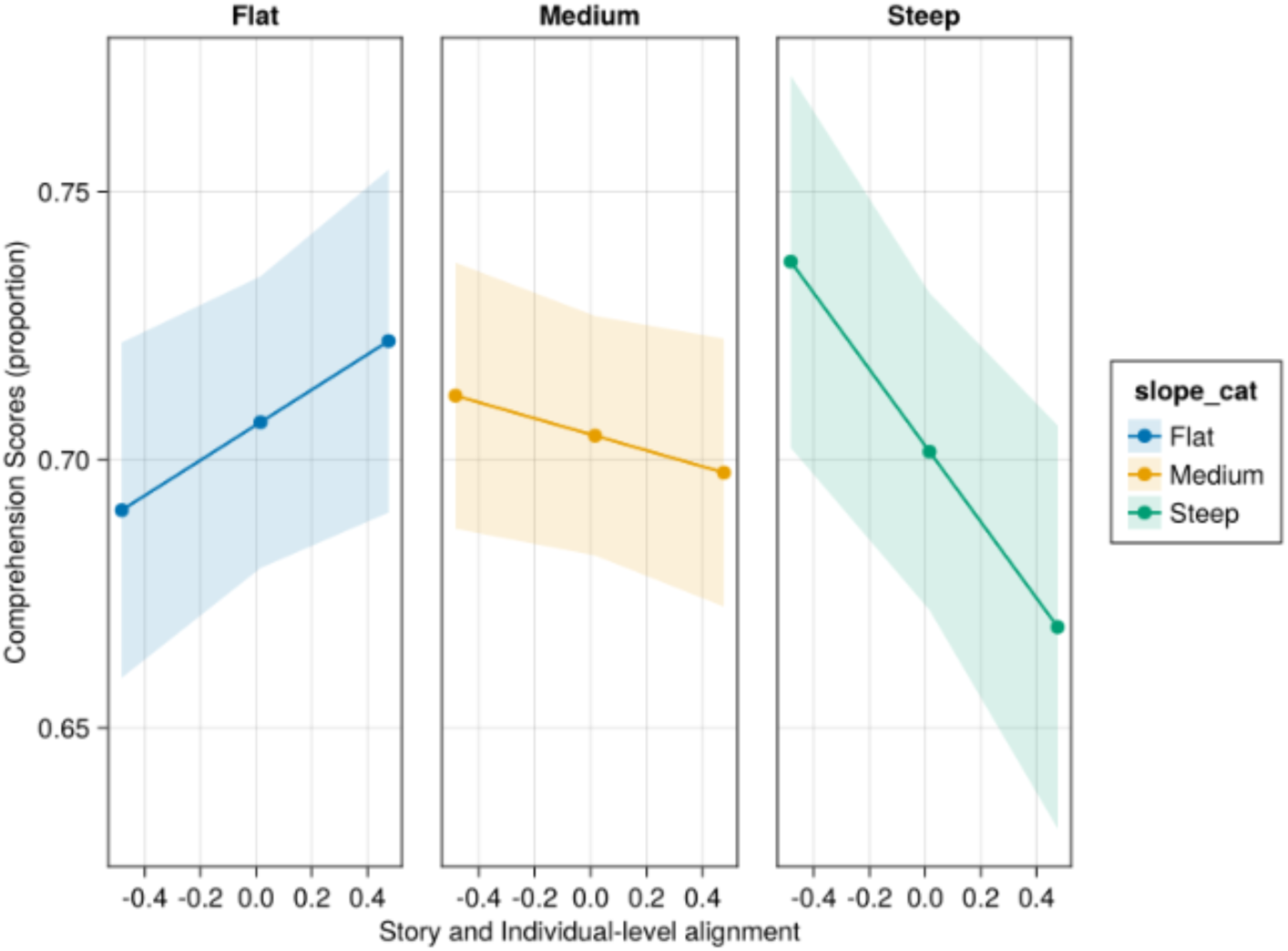
*Plot of interaction between of aperiodic slope and alignment to predict comprehension scores from IM3. The x-axis is story and individual-level alignment, and the y-axis is fitted comprehension scores. The plots are trichotomized for visualisation purposes for three levels of slope. Slope is further visualised by colour with flat, medium and steep slope represented by blue, yellow and green respectively*.

#### Surprisal models

In each ROI, the surprisal models reported main effects of surprisal and surprisal^2^ and significant interaction of these predictors. Table 4 outlines model statistics for significant effects. Figure 14 visualises the main effect of surprisal for the Late Negativity ROI (IM4) and indicates a positive relationship between surprisal and comprehension scores.

**Figure 14:**
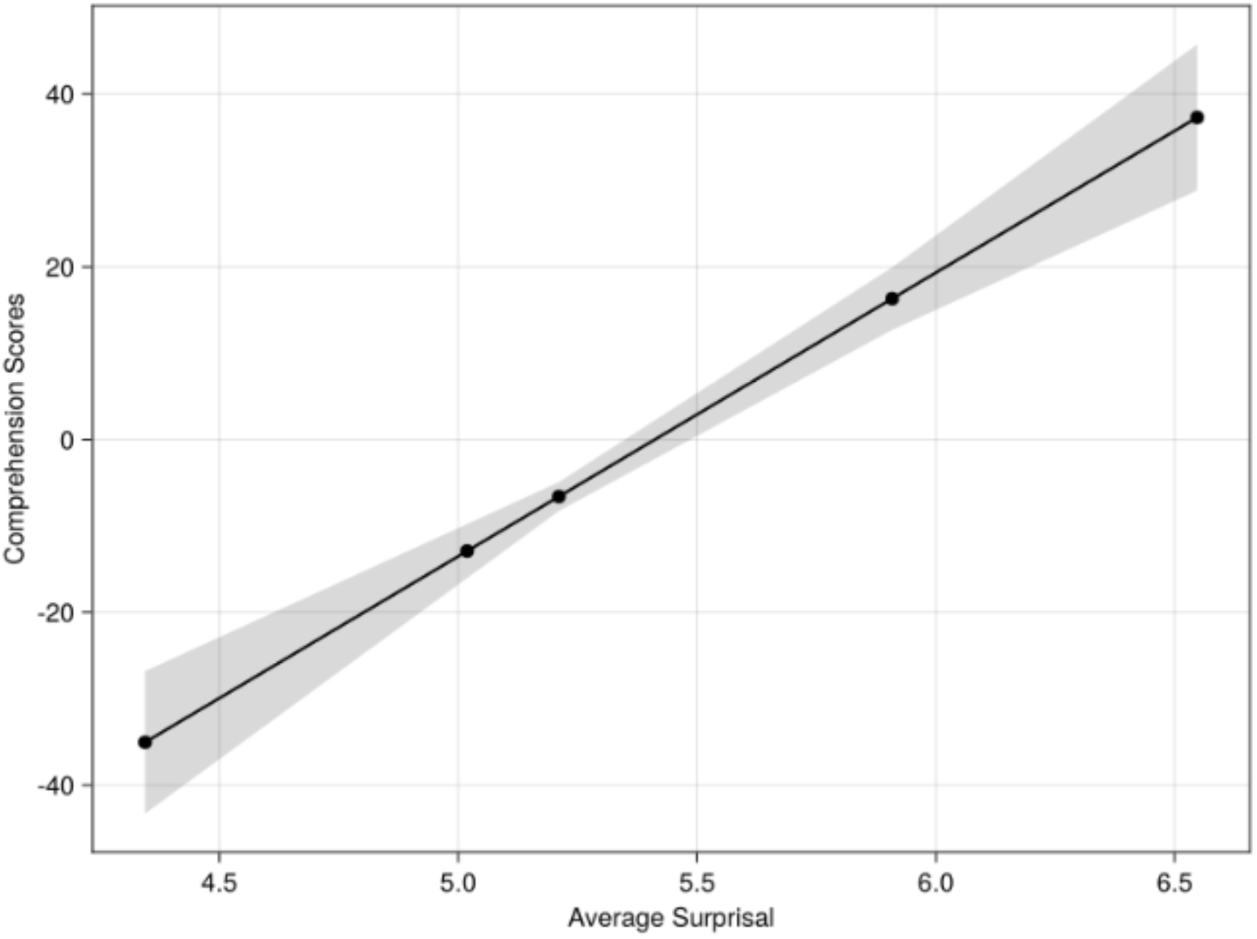
*Significant main effect of story-level surprisal from IM4. The x-axis is story-level surprisal and the y-axis is fitted comprehension scores. The shaded area represents a 95% confidence interval*.

**Table 4:** LMM results for surprisal comprehension models using combined data set

| Model | Window | Significant effect | Estimate | Std. Error | z | p |
| --- | --- | --- | --- | --- | --- | --- |
| LM2 | Traditional N400 | Surprisal | 32.8 | 13.24 | 2.48 | .013 |
|  |  | Surprisal<sup>2</sup> | -6.31 | 2.47 | -2.56 | .011 |
|  |  | Surprisal* | 0.40 | 0.15 | 2.64 | .008 |
|  |  | Surprisal<sup>2</sup> |  |  |  |  |
| LM4 | Late Negativity | Surprisal | 40.27 | 13.97 | 2.88 | .004 |
|  |  | Surprisal<sup>2</sup> | -7.63 | 2.58 | -2.96 | .003 |
|  |  | Surprisal* | 0.47 | 0.15 | 3.03 | .002 |
|  |  | Surprisal<sup>2</sup> |  |  |  |  |

## 4 Discussion

The present study aimed to explore how the interplay between internal and external processing conditions influences the relationship between predictability and language comprehension. Further, the present study aimed to extend the proposal of *Functional Unexpectedness* (Platt, 2025b) to include consideration for modality specific effects and interindividual differences in information processing, as measured by aperiodic slope. It was proposed that an individual’s ability to utilise unexpected information to support language comprehension would be modulated by alignment and aperiodic slope and that we would replicate support for functional unexpectedness, as outlined in Platt (2025a). In an audio-visual analysis, four models were computed, two of which included a surprisal predictor. In these models a significant effect of surprisal was found, demonstrating greater comprehension of more surprising stories. This replicated previous findings in the auditory domain and suggests that more broadly that unexpected information can support understanding of linguistic inputs. Further, in a reanalysis of participants from Platt (2025a) combined with the audio-visual sample reported here, a comprehension model (IM1) including average EEG amplitudes as a predictor demonstrates a significant interaction of aperiodic slope, alignment and EEG amplitudes in predicting comprehension scores. These findings are visualised in Figure 15. An additional model (IM3) in the late negativity ROI, showed that for individuals with a steeper aperiodic slope, lower alignment was related to better comprehension. This model also showed a relationship between higher alignment and improved comprehension for individuals with a flatter aperiodic slope regardless of story-level predictability. From a general comprehension perspective, without considering the role of predictability, this suggests a role of story- and individual-level characteristics in predicting depth of language comprehension. The following sections will further outline this pattern and consider what these findings mean for the account of *Functional Unexpectedness*.

**Figure 15:**
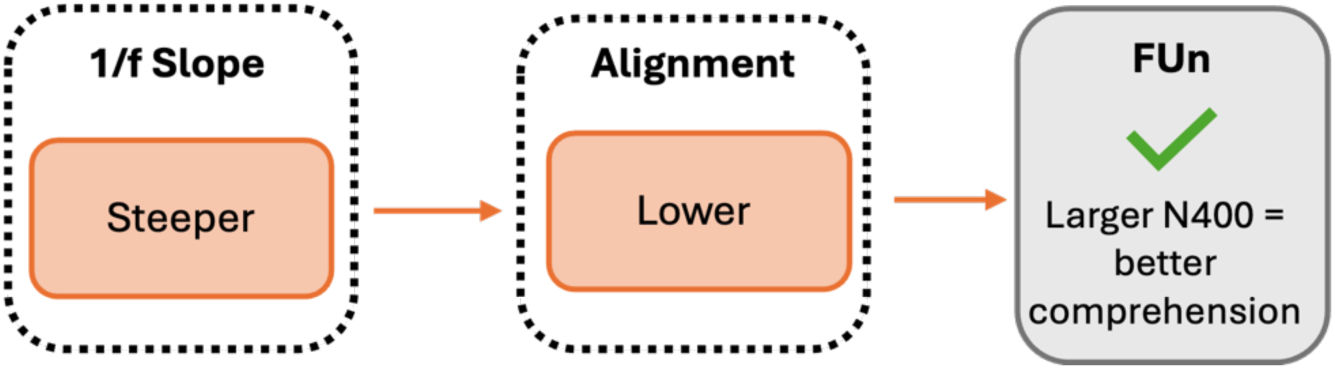
*Processing conditions which lead to Functional Unexpectedness. Findings from the combined analysis N400ROI EEG model (IM1)*.

### 4.1 Processing conditions leading to Functional Unexpectedness

The results from our combined analysis suggest an interplay between aperiodic slope, alignment in predicting comprehension which outlines a series of processing conditions which are related to a functional role of unexpectedness, where greater N400 amplitudes are associated with greater comprehension scores. This finding supports the prediction that the relationship between predictability and language comprehension would be modulated by aperiodic slope and alignment and provides insight into how these characteristics are associated with different behavioural outcomes. It is proposed that the combination of factors, steeper aperiodic slope paired with lower alignment, which leads to functional unexpectedness represents a matching of similarly precision-weighted processing conditions. More specifically it is suggested that both a steeper aperiodic slope and lower alignment weight model updating towards the incoming stimuli and in the case of unexpected information this leads to better comprehension. This suggestion is in line with precision-weighted model updating which outlines that when prior beliefs (existing internal models) are held with higher precision than the incoming stimuli it is less likely that a prediction error will shift the posterior belief towards the unexpected information (Adams et al., 2013; Feldman & Friston, 2010). The present findings further indicate that this likelihood of model updating is modulated by alignment and aperiodic slope. When considering the role of prediction errors in driving model updating, these results suggest that there may be a difference in the ‘gain’ of predictions errors relating to processing conditions. The N400 component has been proposed to reflect an implicit learning signal (Hodapp & Rabovsky, 2021), which reflects the magnitude of model updating in response to new information. Differences in these updating processes have been attributed more generally to overarching individual differences or processing conditions. However, the interplay between alignment and aperiodic slope suggests that processing differences occur across multiple characteristics resulting in processing which can differ based on both internal and external circumstances. Precision-weighting suggests that not all predictions are equal with respect to model updating, and the interplay between alignment and aperiodic slope in the present findings further demonstrates these individual-level weightings and their relationship to comprehension depth.

It was proposed that the novel alignment metric represents the strength of existing models and findings from the present study suggest that this metric may also indicate whether model updating is weighted towards the existing model (high alignment) or the incoming sensory information (lower alignment). A key finding in support of this prediction is the main effect of story- and individual-level alignment in IM1 which showed that lower alignment was associated with greater comprehension scores. If individuals with lower alignment are weighting their model updating towards the incoming signal, they may be more readily integrating story-level information into existing schemas to call on during the comprehension task. This proposal builds on similar suggestions in literature which has explored age-related changes in predictive model adaptation. It has been proposed that older individuals, with a lifetime of experience, hold high precision predictive models – and thus weight predictive processing towards existing models (Moran et al., 2014). As a result, older adults rely on a ‘riskier processing strategy’ (Rayner 2006; Dede 2014) which bases predictions on existing probabilistic expectations, rather than flexibly adapting to local context. Higher alignment individuals may also adopt this ‘risky’ strategy, due to stronger existing schemas weighting processing towards existing models, which does not support better understanding of unexpected information, as unpredictable inputs must be integrated into existing structures.

Aperiodic slope has also been associated with individual differences in model updating (Bornkessel-Schlesewsky, Sharrad, et al., 2022) and age related changes in sensory weighting (Bornkessel-Schlesewsky, Alday, et al., 2022; Moran et al., 2014). Studies have associated aging with a flattening of aperiodic slope (Donoghue et al., 2020; Merkin et al., 2023; Voytek et al., 2015; Waschke et al., 2017) and suggest that this is a reflection that older adults weight model updating more strongly towards existing models (Bornkessel-Schlesewsky, Alday, et al., 2022). Therefore, we suggest that a steeper aperiodic slope gives greater weight to incoming sensory signals. Returning to the role of unexpected stimuli in these relationships, the present study provides evidence that the congruent pairing of these precision-weighted characteristics can provide a pathway to functional unexpectedness. Where high-precision prediction errors are associated with greater model updating, the role of N400 amplitudes suggests that this congruency may support improved comprehension for unexpected stories (as indexed by larger N400 amplitudes). Further, in the case of the opposite congruent pairing (flatter aperiodic slope & higher alignment), where weighting is shifted toward existing models, we see the traditional relationship between predictability and comprehension emerge (see Figure 16 for a visual representation of these weightings). Intriguingly, even in the cases of incongruent pairings (see for example lower alignment & flatter slope in Figure 12) this traditional relationship appears, further highlight the importance of a complementary weighting of both characteristics towards sensory input. Finally, the traditional N400 amplitude and comprehension relationship where less negative amplitudes are associated with better comprehension is evident across almost all other levels of internal and external conditions, with only steeper aperiodic slope and lower alignment leading to a function unexpectedness relationship. This could be evidence that these are the only processing conditions associated with functional unexpectedness, or that this finding is unique to this linguistic context. In favour of the second explanation, the story stimuli presented were written for a general audience to understand and therefore sit towards to middle of a predictability spectrum (extremely predictable – extremely unpredictable). By design, the difference between higher aligned and lower aligned individuals in the processing of more general naturalistic stimuli will not be as extreme compared to presenting stimuli which relate to more specific schemata (e.g., Harry Potter Knowledge; <u>Troyer & Kutas, 2020)</u>. Exploring the role of aperiodic slope and alignment on the comprehension depth of a broader range of naturalistic stimuli, varying in predictability and specificity, would provide insight into the bounds of this functional relationship between lower alignment, steeper slope and depth of comprehension.

**Figure 16:**
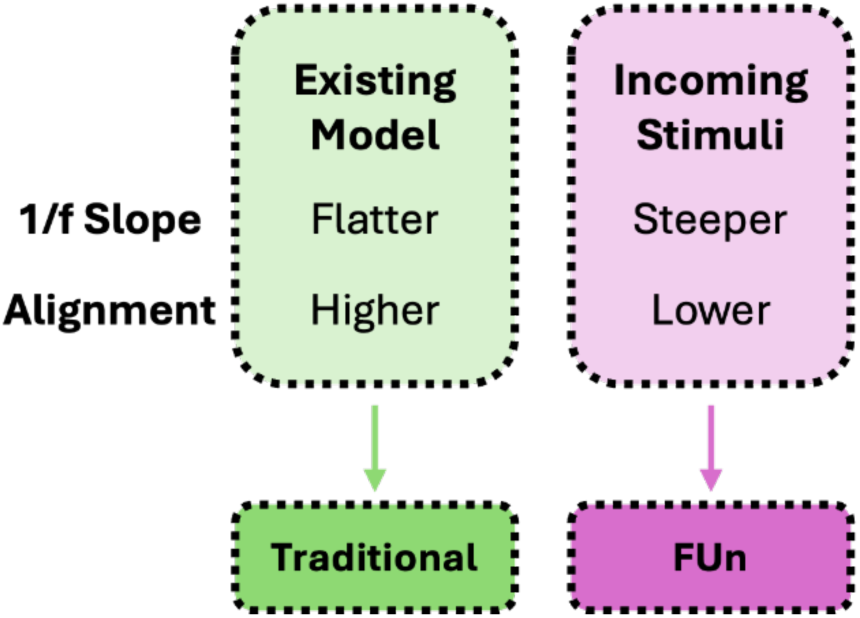
*Proposed weightings of alignment and aperiodic slope characteristics*

### 4.2 Late negativity ROI

The late negativity region of interest (IM3 & IM4) was identified from time and electrode locations where the relationship between word-level average EEG amplitudes and surprisal was significantly correlated. Based on Figure 3 which shows the effect of frequency on EEG amplitudes, it is not surprising that this ROI was identified outside of the traditional N400 ROI. Instead, this data driven approach identified a later negativity ranging from 558ms to 1000ms. This effect was consistent across both modalities, as a comparable latency was identified in Platt (2025a) for the audio-only analysis. This later latency could be associated with the complex processing required for a continuous naturalistic speech stream. Alday et al. (2017), as discussed in context of the word frequency effects above, also demonstrated a ‘shifted N400’ peak at roughly 700ms. In line with the current study, analysis of electrophysiological responses to a 23min naturalistic auditory story in Alday et al. (2017), also showed a frequency effect which peak outside of the traditional 400ms. This finding was attributed to large variations in word length and a note that word-level effect in a naturalistic context reflect the sum of context and other complex interactions. In a non-naturalistic context, Wolff et al. (2008) compared the processing of word order variations for Japanese sentences in an auditory and visual modality. Across both modalities, a consistent posterior negativity between 650-1050ms was present for the processing of object-initial sentences. Wolff et al. (2008) associated this later effect to greater global processing costs for object-initial sentences, which have also been associated with increased error rates and reaction times. This finding in particular may suggest that the complex processing of naturalistic stimuli, and subsequently unexpected information may result in a delayed N400 effect.

There are of course, other language related negativities such as the Mismatch Negativity (MMN) and Left Anterior Negativity (LAN), which could potentially be represented within the ‘late negativity’. However, unlike the effects discussed above these two components appear much earlier in neurophysiological responses. The MMN in particular peaks between 100 and 250ms after a deviant stimulus and is suggest to be elicited by infrequent acoustic events, regardless of focussed attention (Näätänen et al., 1978). Morphosyntactic processing difficulties have been proposed to a elicit an LAN (Gunter et al., 1997). These effects have also been attributed to increased working memory demands in response to morphosyntactic violations (Daneman & Carpenter, 1980; Hasting & Kotz, 2008). However, these later effects are still often only explored prior to 600ms. Suggesting that it may be more plausible to consider the present ‘late negativity’ as a shifted N400 response, in line with Alday et al. (2017). Further research will need to explore the reproducibility of the ‘late-negativity’ in the present analysis and consider its characteristics in comparison with the N400 component and other negativities to provide more insight into the processing it reflects. Nonetheless, this finding motivates the importance of considering how the relationships of interest can inform the analysis process and complement more traditional ERP windows.

### 4.3 Future directions

The current findings identify possible areas for further exploration into the relationship between predictability and language comprehension. Firstly, an important role of alignment has been established in the processing of unexpected information, specially that lower alignment, in conjunction with steeper aperiodic (1/f) slope, is associated with better comprehension of unexpected stories. However, it is unclear what the boundaries of this ‘lower alignment’ processing condition is. An explicit exploration of these boundaries could incorporate the building of a novel schema; in a naturalistic context this might include a unique fictional world created for the experiment. This would provide a timepoint where an individual would be expected to have the lowest level of alignment which could then act as a baseline for increasing alignment as world knowledge is gathered throughout the experiment. A design such as this, would also have the capacity to further consider the role of modality and aperiodic (1/f) activity during this processing. Importantly, findings from this approach would allow for more explicit predictions regarding the level of prior experience necessary to support processing of unexpected stories. Additionally, the alignment metric could be evaluated against an explicit measure of whether individuals have built a new schema during this experiment providing evidence to support the assumption that this measure can act as a proxy for schema strength.

### 4.4 Conclusion

The present study aimed to extend the account of *Functional Unexpectedness* and consider the role of modality and inter-individual differences during the processing of naturalistic language. Across multiple analyses a functional role of surprisal was shown, where more surprising stories were associated with greater comprehension. When considering EEG amplitudes, this relationship was underpinned by a complementary pairing of internal processing conditions. In fact, only when alignment and aperiodic slope were weighted towards the sensory input (i.e., lower alignment & steeper slope) was greater unexpectedness associated with improved comprehension. Outside of this pairing, the more traditional supportive role of increased predictability was demonstrated. Further, an additional EEG model also found that congruent pairings were better for improved comprehension overall. These findings highlight why a functional role of unexpectedness may not have been demonstrated previously in the language processing domain and that this functionality relies on a balance between internal and individual-level processing characteristics. These considerations are essential when considering how individuals may differ in their processing of unexpected information.

## Supplementary Material A: Naturalistic story stimuli examples

### F02 – The new food

I see from the current columns of the daily press that "Professor Plumb, of the University of Chicago, has just invented a highly concentrated form of food. All the essential nutritive elements are put together in the form of pellets, each of which contains from one to two hundred times as much nourishment as an ounce of an ordinary article of diet. These pellets, diluted with water, will form all that is necessary to support life. The professor looks forward confidently to revolutionizing the present food system. "Now this kind of thing may be all very well in its way, but it is going to have its drawbacks as well. In the bright future anticipated by Professor Plumb, we can easily imagine such incidents as the following:

The smiling family were gathered round the hospitable board. The table was plenteously laid with a soup-plate in front of each beaming child, a bucket of hot water before the radiant mother, and at the head of the board the Christmas dinner of the happy home, warmly covered by a thimble and resting on a poker chip. The expectant whispers of the little ones were hushed as the father, rising from his chair, lifted the thimble and disclosed a small pill of concentrated nourishment on the chip before him. Christmas turkey, cranberry sauce, plum pudding, mince pie--it was all there, all jammed into that little pill and only waiting to expand. Then the father with deep reverence, and a devout eye alternating between the pill and heaven, lifted his voice in a benediction. At this moment there was an agonized cry from the mother.

"Oh, Henry, quick! Baby has snatched the pill!" It was too true. Dear little Gustavus Adolphus, the golden-haired baby boy, had grabbed the whole Christmas dinner off the poker chip and bolted it. Three hundred and fifty pounds of concentrated nourishment passed down the oesophagus of the unthinking child. "Clap him on the back!" cried the distracted mother. "Give him water!"

The idea was fatal. The water striking the pill caused it to expand. There was a dull rumbling sound and then, with an awful bang, Gustavus Adolphus exploded into fragments! And when they gathered the little corpse together, the baby lips were parted in a lingering smile that could only be worn by a child who had eaten thirteen Christmas dinners.

### N02 – Colour, spectators to greet 2000 ‘Tour’

Planning is well underway for Strathalbyn and Goolwa’s involvement in the Jacob’s Creek Tour Down Under Cycle Race on January 19. Both towns are making sure colour and excitement will be prominent features for the race which will have international TV coverage. Stage 2 starts at North Adelaide through the Adelaide Hills and into Strathalbyn for the sprint in Dawson Street and then into Goolwa for the race finish in Cadell Street. The towns will welcome the cyclists with their own decoration themes.

Strathalbyn will greet the cyclists with a large painted mural on the hillside at Castle Hill and a town entrance rural theme display leading into a tartan painted road and hundreds of coloured balloons through Dawson Street for the TAB Sprint. Goolwa’s theme will be river and paddle steamers entrance statement across Cadell Street from Moore Streets and colourful flags and sails to the finish line by the Goolwa Hotel.

Businesses and householders are encouraged to decorate their premises along the route in the tour’s colours of Red. Blue and Yellow to be judged by KESAB on the day. There are awards for the Best Decorated Business and Best Decorated House on route in each town and a special award for the Best Dressed Town overall. Entry forms for the Best Dressed Business and House competitions are available from Strathalbyn Tourism Information Centre and at Goolwa from Alexandrina Council Offices. Closing date for entries is Friday, January 14.

Roving entertainers, a Jazz Band, special race events, Sky Dive and a Micro Lite Fly By are just some of the events planned for January 19th 2000 when Goolwa hosts the finish of Stage 2 of the Jacob’s Creek Tour Down Under Cycle Race. Up to 100 of the world’s best cyclists will be taking part in this exciting race which will be enjoyed by an international television audience. The six stage race takes place in South Australia from January 18-23, 2000 and promises to be an even bigger event than the first ’Tour Down Under’ earlier this year.

Pre-race street activities at Goolwa include the SA Veterans & Ladies Cycling Classic Race, Speed Skating, and a Mountain Bike Stunt Team in the children’s park behind Cadell Street where there will also be sponsors’ displays, food and drink stalls, children’s mini fair and face painting. Stuart O’Grady has autographed three bicycle helmets and Tour Down Under T-Shirts for prizes in a competition for under fifteen year olds to be judged on the spot on the day for the best banner or flag brought to cheer the cyclists as they arrive in Goolwa.

### S06 – New research sheds light on how dogs became dogs

At first blush, the emergence of man’s best friend is pretty straightforward. The first dogs descended from wolves in Europe about 14,000 years ago. Then humans domesticated those proto-dogs until the eventual animal known as a "dog" had many of the traits we associated with the animal today. That much of the evolutionary history of the modern dog has been clearly understood. But further research suggests that the European dog is not the ancestor of all our dogs; instead, every modern Western dog hails from a Southeast Asian progenitor lineage. Why? Why did some upstart Southeast Asian lineage triumph, even in Europe, instead of the endemic European one? Turns out, it might have to do with your pet dog’s affinity for Cheetos.

According to research conducted by Ben Sacks from the University of California at Davis and his colleagues, the Southeast Asian dogs prospered because, after they were brought south of the Yangtze River some 6,000 years ago, the dogs were isolated from their wolf forebears. Without that proximity, the Southeast Asian dogs could no longer interbreed with wolves, and thus followed their own evolutionary path. In contrast, northern Asian and European dogs still had contact with, and interbred with, the native wolf populations. Put more clearly, if dogs and wolves interbreed, as they did in Europe, they ended up in an evolutionary cul-de-sac. Isolated from one another, traits that benefited the newly emergent dog lineage flourished.

Another slice of data, published in January in Nature by Erik Axelsson of Uppsala University, suggests that one of the main differences between dogs and wolves is their ability to digest starch. As dogs continued their co-existence with humans - who were, at the same time, mastering agriculture and switching to a grain-based diet - those individuals who could eat starch would be better suited to a domesticated lifestyle than those who had to constantly hunt. Sacks and Axelsson disagree on when that switch took place: Axelsson says that it happened before humans switched from a hunter-gatherer lifestyle to a farming one; Sacks says that the mutation occurred once rice cultivation in Southeast Asia was well underway.

Further work will be required to pinpoint just when the modern dog eventually emerged and to clarify where other canids, such as dingoes, fit into this evolutionary picture. In the interim, sit back and marvel at the process that eventually produced the donut-stealing canine scamps we know and love today.

## Supplementary Material B: Model Output Summaries

**Table SMB-1:** Model outputs for AM1

| Effect | Coef | Std.<br>error | z | p |
| --- | --- | --- | --- | --- |
| Intercept | 0.17 | 0.09 | 1.82 | .069 |
| ps | 1.47 | 0.17 | 8.87 | <.001 |
| word (centered: 222.55 scaled: 129.02) | -0.01 | 0.05 | -0.21 | .836 |
| lfg (centered: 5.42 scaled: 1.49) | -0.01 | 0.07 | -0.16 | .871 |
| ls_sum(centered: 5.49 scaled: 4.4) | 0.10 | 0.06 | 1.65 | .099 |
| dur (centered: 0.33 scaled: 0.2) | -0.17 | 0.03 | -6.22 | <.001 |
| mean freq(centered: 227.35 scaled: 46.74) | 0.14 | 0.01 | 11.47 | <.001 |
| mean_int(centered: 58.42 scaled: 8.02) | 0.04 | 0.01 | 3.79 | <.001 |
| fra pre(centered: 4.97682497e6 scaled: 8.03645744e6) | 0.23 | 0.02 | 13.25 | <.001 |
| dur_pre(centered: 0.32 scaled: 0.2) | 0.03 | 0.01 | 2.89 | .004 |
| ps & word (centered: 222.55 scaled: 129.02) | -0.09 | 0.01 | -7.38 | <.001 |
| ps & lfg(centered: 5.42 scaled: 1.49) | 0.03 | 0.05 | 0.56 | .523 |
| word (centered: 222.55 scaled: 129.02) & lfg (centered: 5.42 scaled: 1.49) | -0.05 | 0.05 | -0.89 | .374 |
| ps & ls_sum(centered: 5.49 scaled: 4.4) | 0.01 | 0.02 | 0.41 | .681 |
| word (centered: 222.55 scaled: 129.02) & ls_sum(centered: 5.49 scaled: 4.4) | -0.07 | 0.02 | -3.53 | <.001 |
| lfg(centered: 5.42 scaled: 1.49) & ls_sum(centered: 5.49 scaled: 4.4) | 0.02 | 0.05 | 0.40 | .689 |
| ps & word (centered: 222.55 scaled: 129.02) & lfg (centered: 5.42 scaled: 1.49) | 0.08 | 0.01 | 6.38 | <.001 |
| ps & word (centered: 222.55 scaled: 129.02) & ls_sum(centered: 5.49 scaled: 4.4) | 0.02 | 0.01 | 1.27 | .204 |
| ps & lfg(centered: 5.42 scaled: 1.49) & ls_sum(centered: 5.49 scaled: 4.4) | 0.04 | 0.01 | 2.81 | .005 |
| word (centered: 222.55 scaled: 129.02) & lfg (centered: 5.42 scaled: 1.49) & ls_sum(centered: 5.49 scaled: 4.4) | 0.05 | 0.02 | 2.37 | .018 |
| ps & word (centered: 222.55 scaled: 129.02) & lfg (centered: 5.42 scaled: 1.49) & ls_sum(centered: 5.49 scaled: 4.4) | 0.02 | 0.01 | 1.53 | .126 |

**Table SMB-2:**
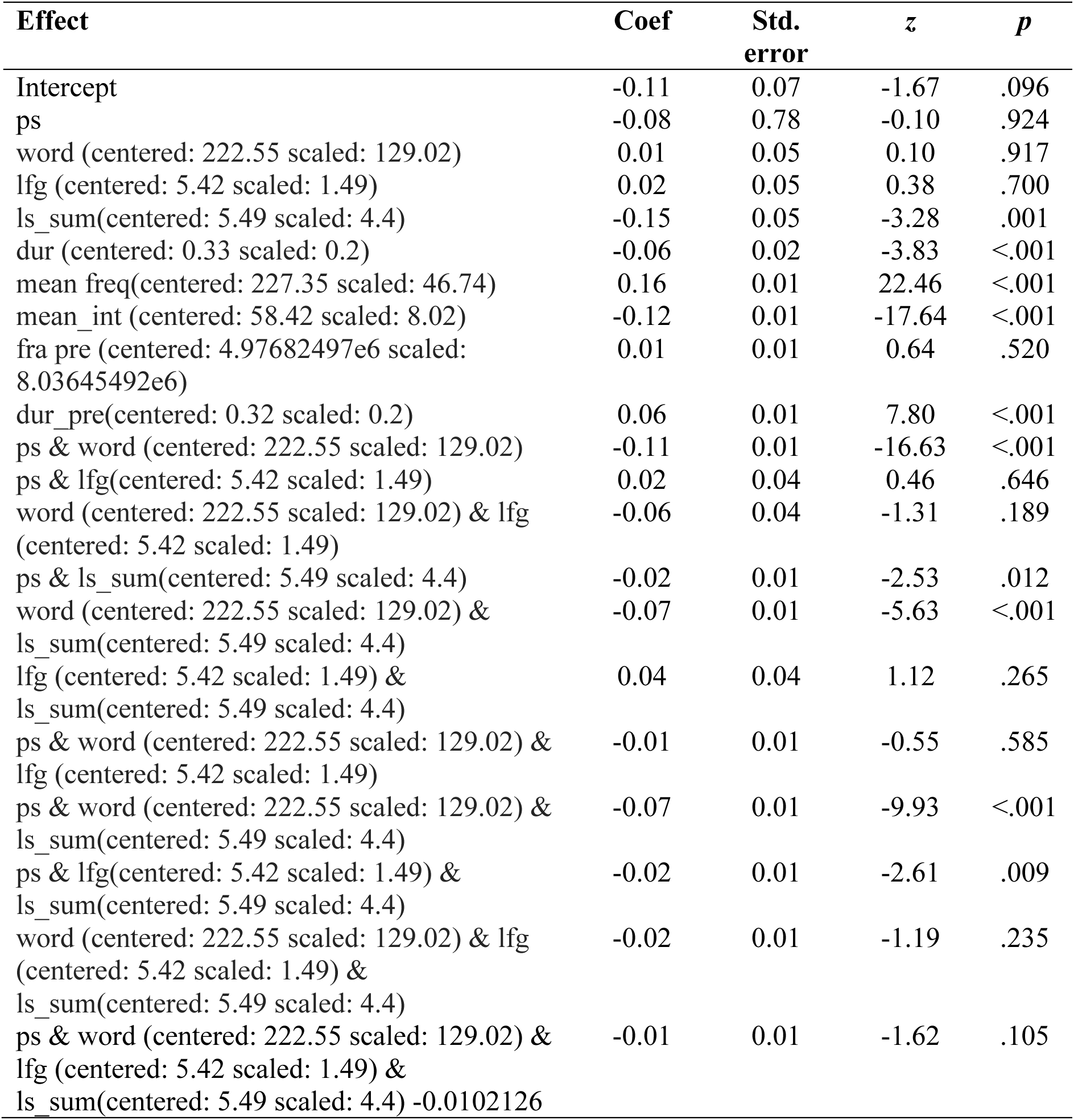
Model outputs for AM2

| Effect | Coef | Std.<br>error | z | p |
| --- | --- | --- | --- | --- |
| Intercept | -0.11 | 0.07 | -1.67 | .096 |
| ps | -0.08 | 0.78 | -0.10 | .924 |
| word (centered: 222.55 scaled: 129.02) | 0.01 | 0.05 | 0.10 | .917 |
| lfg (centered: 5.42 scaled: 1.49) | 0.02 | 0.05 | 0.38 | .700 |
| ls_sum(centered: 5.49 scaled: 4.4) | -0.15 | 0.05 | -3.28 | .001 |
| dur (centered: 0.33 scaled: 0.2) | -0.06 | 0.02 | -3.83 | <.001 |
| mean freq(centered: 227.35 scaled: 46.74) | 0.16 | 0.01 | 22.46 | <.001 |
| mean_int (centered: 58.42 scaled: 8.02) | -0.12 | 0.01 | -17.64 | <.001 |
| fra pre (centered: 4.97682497e6 scaled: 8.03645492e6) | 0.01 | 0.01 | 0.64 | .520 |
| dur_pre(centered: 0.32 scaled: 0.2) | 0.06 | 0.01 | 7.80 | <.001 |
| ps & word (centered: 222.55 scaled: 129.02) | -0.11 | 0.01 | -16.63 | <.001 |
| ps & lfg(centered: 5.42 scaled: 1.49) | 0.02 | 0.04 | 0.46 | .646 |
| word (centered: 222.55 scaled: 129.02) & lfg (centered: 5.42 scaled: 1.49) | -0.06 | 0.04 | -1.31 | .189 |
| ps & ls_sum(centered: 5.49 scaled: 4.4) | -0.02 | 0.01 | -2.53 | .012 |
| word (centered: 222.55 scaled: 129.02) & ls_sum(centered: 5.49 scaled: 4.4) | -0.07 | 0.01 | -5.63 | <.001 |
| lfg (centered: 5.42 scaled: 1.49) & ls_sum(centered: 5.49 scaled: 4.4) | 0.04 | 0.04 | 1.12 | .265 |
| ps & word (centered: 222.55 scaled: 129.02) & lfg (centered: 5.42 scaled: 1.49) | -0.01 | 0.01 | -0.55 | .585 |
| ps & word (centered: 222.55 scaled: 129.02) & ls_sum(centered: 5.49 scaled: 4.4) | -0.07 | 0.01 | -9.93 | <.001 |
| ps & lfg(centered: 5.42 scaled: 1.49) & ls_sum(centered: 5.49 scaled: 4.4) | -0.02 | 0.01 | -2.61 | .009 |
| word (centered: 222.55 scaled: 129.02) & lfg (centered: 5.42 scaled: 1.49) & ls_sum(centered: 5.49 scaled: 4.4) | -0.02 | 0.01 | -1.19 | .235 |
| ps & word (centered: 222.55 scaled: 129.02) & lfg (centered: 5.42 scaled: 1.49) & ls_sum(centered: 5.49 scaled: 4.4) -0.0102126 | -0.01 | 0.01 | -1.62 | .105 |

**Table SMB-3:** Model outputs for CM1

| Effect | Coef | Std.<br>error | z | p |
| --- | --- | --- | --- | --- |
| Intercept | -0.86 | 0.69 | -1.25 | .213 |
| mean_sentence_length | -0.03 | 0.01 | -2.88 | .004 |
| mean_word_dur | 5.37 | 1.75 | 3.06 | .003 |
| mean_unigram_freq (centered: 4.87976251e6<br>scaled: 637881.49) | 0.02 | 0.03 | 0.66 | .512 |
| idea_density | 0.63 | 0.65 | 0.96 | .335 |
| genre: newspaper | 0.13 | 0.06 | 2.09 | .036 |
| genre: science | -0.18 | 0.06 | -2.90 | .004 |
| alignment_value | 0.03 | 0.04 | 0.75 | .454 |
| mean_mean | 0.03 | 0.03 | 0.97 | .331 |
| mean_mean2 | 0.01 | 0.02 | 0.50 | .620 |
| alignment_value & mean_mean | 0.02 | 0.06 | 0.26 | .791 |
| alignment_value & mean_mean2 | -0.04 | 0.03 | -1.25 | .211 |
| mean_mean & mean_mean2 | -6.90 | 0.02 | -0.00 | .996 |
| alignment_value & mean_mean &<br>mean_mean2 | 0.01 | 0.02 | 0.54 | .591 |

**Table SMB-4:** Model outputs for CM2

| Effect | Coef | Std.<br>error | z | p |
| --- | --- | --- | --- | --- |
| Intercept | -39.95 | 9.44 | -4.23 | <.001 |
| mean_sentence_length | -0.09 | 0.01 | -9.24 | <.001 |
| mean_word_dur | 9.37 | 1.20 | 7.78 | <.001 |
| mean_unigram_freq (centered: 4.87976251e6<br>scaled: 637881.49) | 0.13 | 0.02 | 5.55 | <.001 |
| idea_density | 3.80 | 0.55 | 6.93 | <.001 |
| genre: newspaper | 0.27 | 0.04 | 7.68 | <.001 |
| genre: science | -0.26 | 0.04 | -6.30 | <.001 |
| alignment_value | 9.10 | 15.96 | 0.57 | .568 |
| ls_sum_mean_fixed | 21.81 | 5.23 | 4.17 | <.001 |
| ls_sum_mean2_fixed | -4.27 | 0.98 | -4.35 | <.001 |
| alignment_value & ls_sum_mean_fixed | -4.72 | 9.00 | -0.52 | .600 |
| alignment_value & ls_sum_mean2_fixed | 0.81 | 1.68 | 0.48 | .630 |
| ls_sum_mean_fixed & ls_sum_mean2_fixed | 0.28 | 0.06 | 4.53 | <.001 |
| alignment_value & ls_sum_mean_fixed &<br>ls_sum_mean2_fixed | -0.05 | 0.10 | -0.44 | .658 |

**Table SMB-5:** Model outputs for CM3

| Effect | Coef | Std.<br>error | z | p |
| --- | --- | --- | --- | --- |
| (Intercept) | -0.88 | 0.66 | -1.32 | .187 |
| mean_sentence_length | -0.04 | 0.01 | -2.95 | .003 |
| mean_word_dur | 5.38 | 1.70 | 3.17 | .002 |
| mean_unigram_freq(centered: 4.87976251e6<br>scaled: 637881.49) | 0.02 | 0.02 | 0.75 | .453 |
| idea_density | 0.68 | 0.64 | 1.06 | .288 |
| genre: newspaper | 0.13 | 0.06 | 2.18 | .029 |
| genre: science | -0.18 | 0.06 | -3.03 | .003 |
| alignment_value | -0.01 | 0.04 | -0.13 | .900 |
| mean_mean | 0.01 | 0.04 | 0.18 | .858 |
| mean_mean2 | 0.02 | 0.04 | 0.55 | .584 |
| alignment_value & mean_mean | 0.02 | 0.09 | 0.27 | .790 |
| alignment_value & mean_mean2 | -0.04 | 0.07 | -0.52 | .602 |
| mean_mean & mean_mean2 | 0.00 | 0.03 | 0.10 | .924 |
| alignment_value & mean_mean &<br>mean_mean2 | 0.01 | 0.06 | 0.09 | .928 |

**Table SMB-6:**
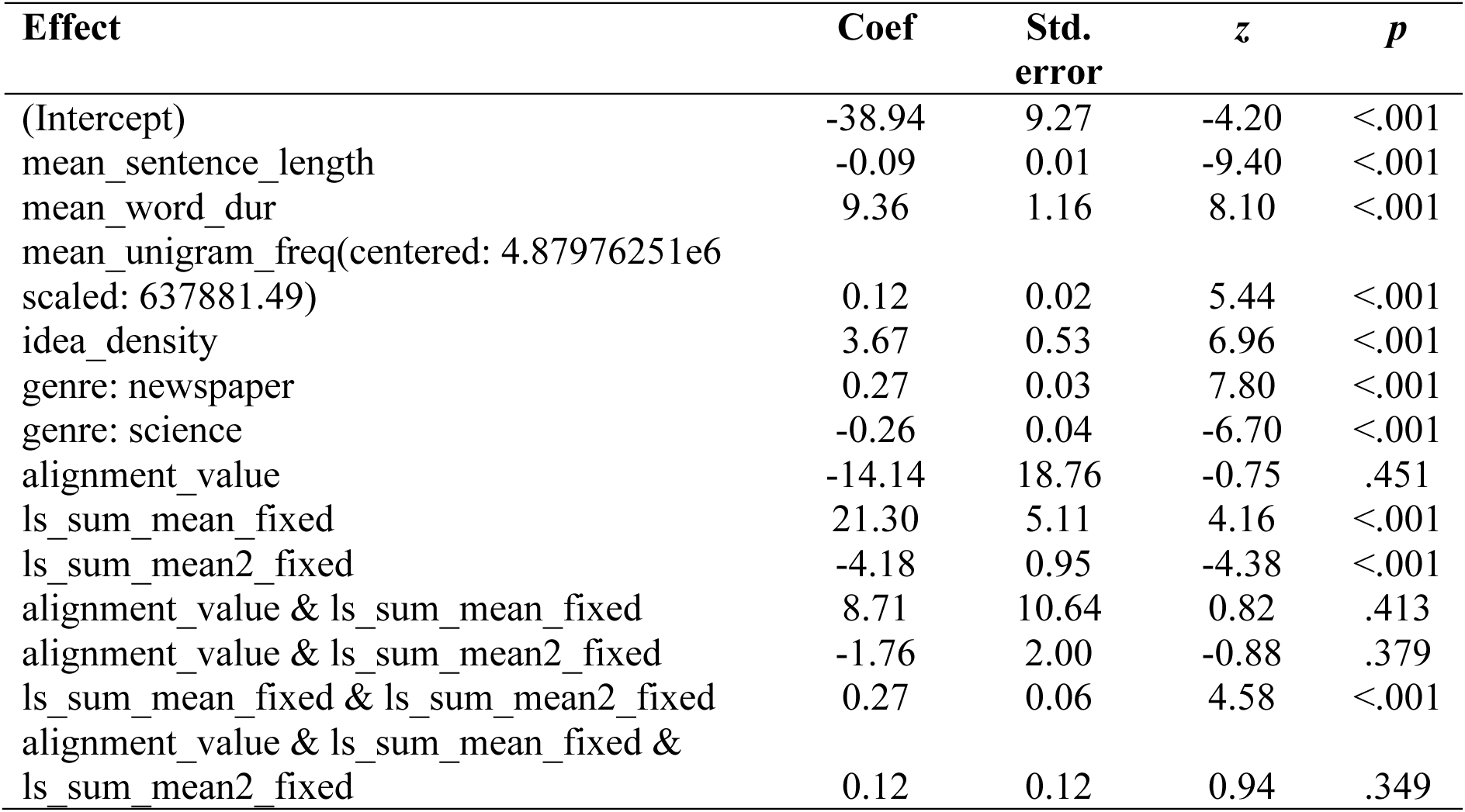
Model outputs for CM4

| Effect | Coef | Std.<br>error | z | p |
| --- | --- | --- | --- | --- |
| (Intercept) | -38.94 | 9.27 | -4.20 | <.001 |
| mean_sentence_length | -0.09 | 0.01 | -9.40 | <.001 |
| mean_word_dur | 9.36 | 1.16 | 8.10 | <.001 |
| mean_unigram_freq(centered: 4.87976251e6<br>scaled: 637881.49) | 0.12 | 0.02 | 5.44 | <.001 |
| idea_density | 3.67 | 0.53 | 6.96 | <.001 |
| genre: newspaper | 0.27 | 0.03 | 7.80 | <.001 |
| genre: science | -0.26 | 0.04 | -6.70 | <.001 |
| alignment_value | -14.14 | 18.76 | -0.75 | .451 |
| ls_sum_mean_fixed | 21.30 | 5.11 | 4.16 | <.001 |
| ls_sum_mean2_fixed | -4.18 | 0.95 | -4.38 | <.001 |
| alignment_value & ls_sum_mean_fixed | 8.71 | 10.64 | 0.82 | .413 |
| alignment_value & ls_sum_mean2_fixed | -1.76 | 2.00 | -0.88 | .379 |
| ls_sum_mean_fixed & ls_sum_mean2_fixed | 0.27 | 0.06 | 4.58 | <.001 |
| alignment_value & ls_sum_mean_fixed &<br>ls_sum_mean2_fixed | 0.12 | 0.12 | 0.94 | .349 |

**Table SMB-7:**
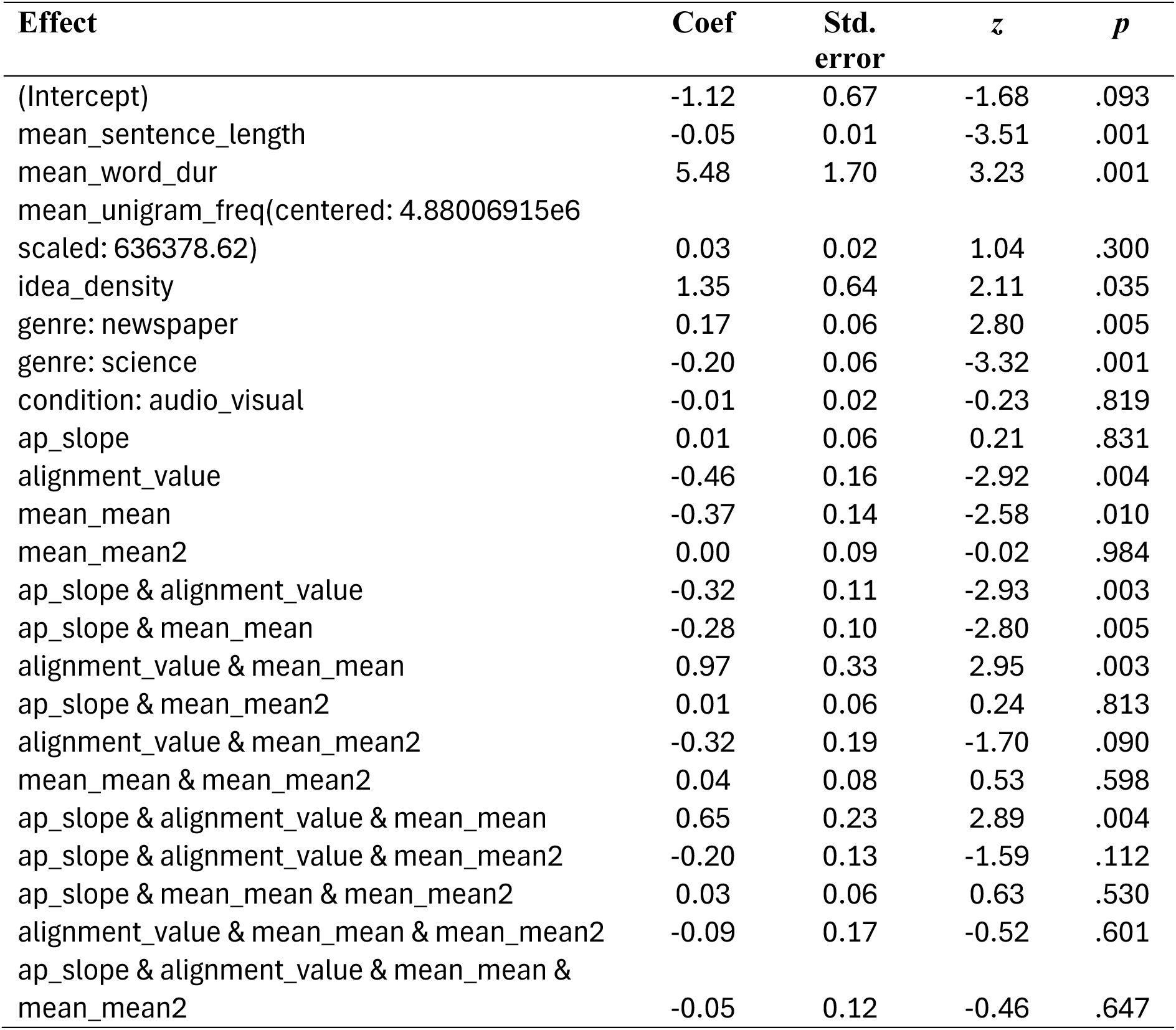
Model outputs for IM1

**Table SMB-8:** Model outputs for IM2

| Effect | Coef | Std.<br>error | z | p |
| --- | --- | --- | --- | --- |
| (Intercept) | -59.90 | 23.62 | -2.54 | .011 |
| mean_sentence_length | -0.10 | 0.01 | -8.37 | <.001 |
| mean_word_dur | 9.27 | 1.40 | 6.61 | <.001 |
| mean_unigram_freq(centered: 4.88006915e6<br>scaled: 636378.62) | 0.13 | 0.03 | 4.94 | <.001 |
| idea_density | 4.30 | 0.64 | 6.70 | <.001 |
| genre: newspaper | 0.30 | 0.04 | 7.30 | <.001 |
| genre: science | -0.28 | 0.05 | -5.78 | <.001 |
| condition: audio_visual | 0.00 | 0.02 | -0.08 | .936 |
| ap_slope | -13.93 | 15.32 | -0.91 | .363 |
| alignment_value | 12.37 | 84.07 | 0.15 | .883 |
| ls_sum_mean_fixed | 32.83 | 13.24 | 2.48 | .013 |
| ls_sum_mean2_fixed | -6.31 | 2.47 | -2.56 | .011 |
| ap_slope & alignment_value | 4.88 | 57.86 | 0.08 | .933 |
| ap_slope & ls_sum_mean_fixed | 7.82 | 8.62 | 0.91 | .364 |
| alignment_value & ls_sum_mean_fixed | -1.99 | 47.60 | -0.04 | .967 |
| ap_slope & ls_sum_mean2_fixed | -1.47 | 1.60 | -0.91 | .361 |
| alignment_value & ls_sum_mean2_fixed | -0.56 | 8.93 | -0.06 | .950 |
| ls_sum_mean_fixed & ls_sum_mean2_fixed | 0.40 | 0.15 | 2.64 | .008 |
| ap_slope & alignment_value &<br>ls_sum_mean_fixed | 0.62 | 32.74 | 0.02 | .985 |
| ap_slope & alignment_value &<br>ls_sum_mean2_fixed | -0.75 | 6.14 | -0.12 | .903 |
| ap_slope & ls_sum_mean_fixed &<br>ls_sum_mean2_fixed | 0.09 | 0.10 | 0.92 | .356 |
| alignment_value & ls_sum_mean_fixed &<br>ls_sum_mean2_fixed | 0.09 | 0.55 | 0.16 | .871 |
| ap_slope & alignment_value &<br>ls_sum_mean_fixed & ls_sum_mean2_fixed | 0.08 | 0.38 | 0.22 | .825 |

**Table SMB-9:** Model outputs for IM3

| Effect | Coef | Std.<br>error | z | p |
| --- | --- | --- | --- | --- |
| (Intercept) | -1.29 | 0.67 | -1.92 | .054 |
| mean_sentence_length | -0.04 | 0.01 | -3.42 | .001 |
| mean_word_dur | 5.48 | 1.70 | 3.23 | .001 |
| mean_unigram_freq(centered: 4.88006915e6<br>scaled: 636378.62) | 0.02 | 0.02 | 1.02 | .308 |
| idea_density | 1.33 | 0.64 | 2.09 | .037 |
| genre: newspaper | 0.17 | 0.06 | 2.76 | .006 |
| genre: science | -0.20 | 0.06 | -3.29 | .001 |
| condition: audio_visual | 0.00 | 0.02 | -0.10 | .922 |
| ap_slope | -0.10 | 0.06 | -1.70 | .089 |
| alignment_value | -0.46 | 0.17 | -2.79 | .005 |
| mean_mean | -0.09 | 0.18 | -0.52 | .606 |
| mean_mean2 | 0.54 | 0.25 | 2.15 | .032 |
| ap_slope & alignment_value | -0.33 | 0.12 | -2.81 | .005 |
| ap_slope & mean_mean | -0.05 | 0.12 | -0.43 | .669 |
| alignment_value & mean_mean | -0.32 | 0.40 | -0.80 | .427 |
| ap_slope & mean_mean2 | 0.38 | 0.17 | 2.31 | .021 |
| alignment_value & mean_mean2 | 0.72 | 0.47 | 1.53 | .126 |
| mean_mean & mean_mean2 | 0.25 | 0.23 | 1.08 | .281 |
| ap_slope & alignment_value & mean_mean | -0.24 | 0.28 | -0.85 | .393 |
| ap_slope & alignment_value & mean_mean2 | 0.53 | 0.31 | 1.68 | .094 |
| ap_slope & mean_mean & mean_mean2 | 0.17 | 0.15 | 1.12 | .262 |
| alignment_value & mean_mean & mean_mean2 | 0.47 | 0.33 | 1.45 | .148 |
| ap_slope & alignment_value & mean_mean &<br>mean_mean2 | 0.33 | 0.22 | 1.52 | .130 |

**Table SMB-10:** Model outputs for IM4

| Effect | Coef | Std.<br>error | z | p |
| --- | --- | --- | --- | --- |
| (Intercept) | -73.75 | 25.14 | -2.93 | .003 |
| mean_sentence_length | -0.09 | 0.01 | -8.90 | <.001 |
| mean_word_dur | 9.26 | 1.31 | 7.06 | <.001 |
| mean_unigram_freq(centered: 4.88006915e6<br>scaled: 636378.62) | 0.13 | 0.02 | 5.15 | <.001 |
| idea_density | 4.29 | 0.60 | 7.18 | <.001 |
| genre: newspaper | 0.30 | 0.04 | 7.81 | <.001 |
| genre: science | -0.28 | 0.05 | -6.27 | <.001 |
| condition: audio_visual | 0.00 | 0.02 | 0.02 | .987 |
| ap_slope | -21.61 | 16.67 | -1.30 | .195 |
| alignment_value | 98.79 | 78.21 | 1.26 | .207 |
| ls_sum_mean_fixed | 40.27 | 13.97 | 2.88 | .004 |
| ls_sum_mean2_fixed | -7.63 | 2.58 | -2.96 | .003 |
| ap_slope & alignment_value | 83.94 | 55.49 | 1.51 | .130 |
| ap_slope & ls_sum_mean_fixed | 11.86 | 9.27 | 1.28 | .201 |
| alignment_value & ls_sum_mean_fixed | -55.71 | 44.42 | -1.25 | .210 |
| ap_slope & ls_sum_mean2_fixed | -2.16 | 1.71 | -1.27 | .206 |
| alignment_value & ls_sum_mean2_fixed | 10.30 | 8.35 | 1.23 | .217 |
| ls_sum_mean_fixed & ls_sum_mean2_fixed | 0.48 | 0.16 | 3.03 | .002 |
| ap_slope & alignment_value &<br>ls_sum_mean_fixed | -47.97 | 31.55 | -1.52 | .128 |
| ap_slope & alignment_value &<br>ls_sum_mean2_fixed | 9.00 | 5.94 | 1.52 | .129 |
| ap_slope & ls_sum_mean_fixed &<br>ls_sum_mean2_fixed | 0.13 | 0.10 | 1.25 | .211 |
| alignment_value & ls_sum_mean_fixed &<br>ls_sum_mean2_fixed | -0.63 | 0.52 | -1.21 | .227 |
| ap_slope & alignment_value &<br>ls_sum_mean_fixed & ls_sum_mean2_fixed | -0.56 | 0.37 | -1.51 | .132 |

